# Limited inhibition of multiple nodes in a driver network blocks metastasis

**DOI:** 10.1101/2020.06.05.137117

**Authors:** Ali E. Yesilkanal, Dongbo Yang, Payal Tiwari, Alan U. Sabino, Jiyoung Lee, Xiao-He Xie, Siqi Sun, Christopher Dann, Ethan Steinberg, Timothy Stuhlmiller, Casey Frankenberger, Elizabeth Goldsmith, Gary L. Johnson, Alexandre F. Ramos, Marsha R. Rosner

## Abstract

Metastasis suppression by high-dose, multi-drug targeting is largely unsuccessful due to network heterogeneity and compensatory network activation. Here we show that targeting driver network signaling capacity by limited inhibition of its core pathways is a more effective anti-metastatic strategy. This principle underlies the action of a physiological metastasis suppressor, Raf Kinase Inhibitory Protein, which moderately decreases stress-regulated MAP kinase network activity, reducing the output to metastatic transcription factor BACH1 and motility-related target genes. We developed a low-dose four-drug mimic that blocks metastatic colonization in mouse breast cancer models and increases survival. Experiments and network flow modeling show: 1) limited inhibition of multiple pathways is required to overcome variation in MAPK network topology and suppress signaling output across heterogeneous tumor cells, and 2) restricting inhibition of individual kinases dissipates surplus signal, preventing threshold activation of compensatory kinase networks. This low-dose multi-drug approach to decrease signaling capacity of driver networks represents a transformative, clinically-relevant strategy for anti-metastatic treatment.

---

Cancer is a complex disease marked by heterogeneity. For solid tumors, metastatic dissemination of sub-populations of tumor cells throughout the body is primarily responsible for lethality^1^. Metastasis is characterized by many distinct biological processes such as tumor cell invasion, transport in vessels, and colonization at distant sites that involve significant cellular stress. Metastatic progression is further complicated by dynamic changes in tumors that undergo evolutionary change in response to cells and stresses within the microenvironment.

Previous approaches to treating metastatic disease have largely been ineffective at preventing resistance or recurrence due to cellular heterogeneity and robust compensatory mechanisms. Targeting individual metastatic pathways at maximum tolerated doses using single or multiple anti-cancer agents can activate compensatory pathways that eventually overcome treatment^2–4^. Even high dose combination therapies that target single kinases across multiple networks can be toxic and are insufficient to enable long-term survival^^5–7^^. The commonality between signaling pathways in both normal and tumor cells has also limited the efficacy of most therapeutic strategies. Therefore, novel strategies for suppressing metastasis are needed, particularly for cancers such as triple negative breast cancer (ER^−^, PR^−^, HER2^low^; TNBC) that lack effective targeted therapy.

An alternative approach to suppressing metastasis employs a phenomenological framework built upon understanding the action of physiological metastasis regulators. Biological metastasis suppressors are proteins that inhibit various steps of metastasis and are lost or silenced in metastatic tumors^8^. To date, approximately 100 metastasis suppressors have been identified, many of which also inhibit tumor growth^8^. Interestingly, several of these metastasis suppressors are kinases or proteins that modify signaling cascades and provide insight into metastatic signaling mechanisms. One of these, Raf Kinase Inhibitory Protein (RKIP; PEBP1), is a regulator of Raf kinase activity that is deleted or lost in virtually all metastatic solid tumors (reviewed by Yesilkanal and Rosner, 2018). Reintroducing RKIP to metastatic TNBC cells blocks invasion of cells *in vitro* and inhibits intravasation and metastasis of tumor cells *in vivo*^9^. Numerous studies have shown that loss of RKIP protein expression is associated with poor outcome in a variety of tumors including breast, prostate, and melanoma^10^. Furthermore, expression of RKIP in preclinical models enhances the response to chemotherapy as well as radiation suggesting that RKIP can potentiate therapeutic efficacy^11–13^. Therefore, RKIP provides a powerful model system for developing new anti-metastatic therapies based on the mechanism by which RKIP modulates signaling network dynamics and prevents metastatic transformation.

In the present study, we utilized the action of RKIP as a conceptual framework for a new strategy to target metastasis. Metastatic suppression is achieved by restricting but not eliminating the activities of multiple kinases in a driver signaling network, stress MAP kinases (MAPKs). The molecular output of the network, the transcription factor BACH1, drives clinically relevant metastatic motility genes. We developed a 4-drug mimic of RKIP used at sub-therapeutic doses that inhibits network output, reduces metastasis, and improves survival *in vivo*. Modeling of different MAPK network topologies provides a rationale for this multi-drug anti-metastatic strategy that reduces signaling flow at multiple rather than single nodes and prevents activation of compensatory signaling pathways. These findings challenge the current approaches to drug treatment and suggest an alternative strategy for controlling metastatic disease in breast and potentially other cancers.

## RESULTS

### RKIP regulates a clinically relevant set of motility-related genes driven by the pro-metastatic transcription factor BACH1

To characterize the mechanism by which RKIP suppresses metastasis, we first analyzed gene expression data from breast cancer patient samples in The Cancer Genome Atlas (TCGA) study. Our analysis revealed that RKIP expression negatively correlated with genes involved in cell motility (cell leading edge, cell migration, focal adhesion) and kinase-mediated signaling (regulation of GTPase activity, phosphotransferase activity) (Fig. 1A). Among the genes most inversely correlated with RKIP was BACH1 (BTB and CNC homology 1), a pro-metastatic, basic leucine zipper transcription factor that is post-translationally inhibited by RKIP via let7 (Fig. 1A)^9, 14^. To test direct regulation of these motility genes by RKIP, we performed RNA sequencing (RNA-seq) of transcripts in control versus RKIP-expressing xenograft tumors of BM1, a bone-tropic, Ras/B-Raf mutant TNBC cell line derived from MDA-MB-231 (Supplementary Fig. 1A,B)^15^. Over 70 of the motility genes as well as BACH1 that inversely correlate with RKIP in patients were also downregulated by RKIP in xenograft tumors, suggesting these genes are transcriptionally regulated by RKIP in TNBC (Fig. 1B,C and Supplementary Fig. 1C). We validated the differential expression of 15 motility genes previously implicated in metastasis^16–24^, using BM1 or MB436 cells expressing wildtype or a more robust version of RKIP (S153E mutant) (Supplementary Fig. 1D)^9^.

**Fig. 1:**
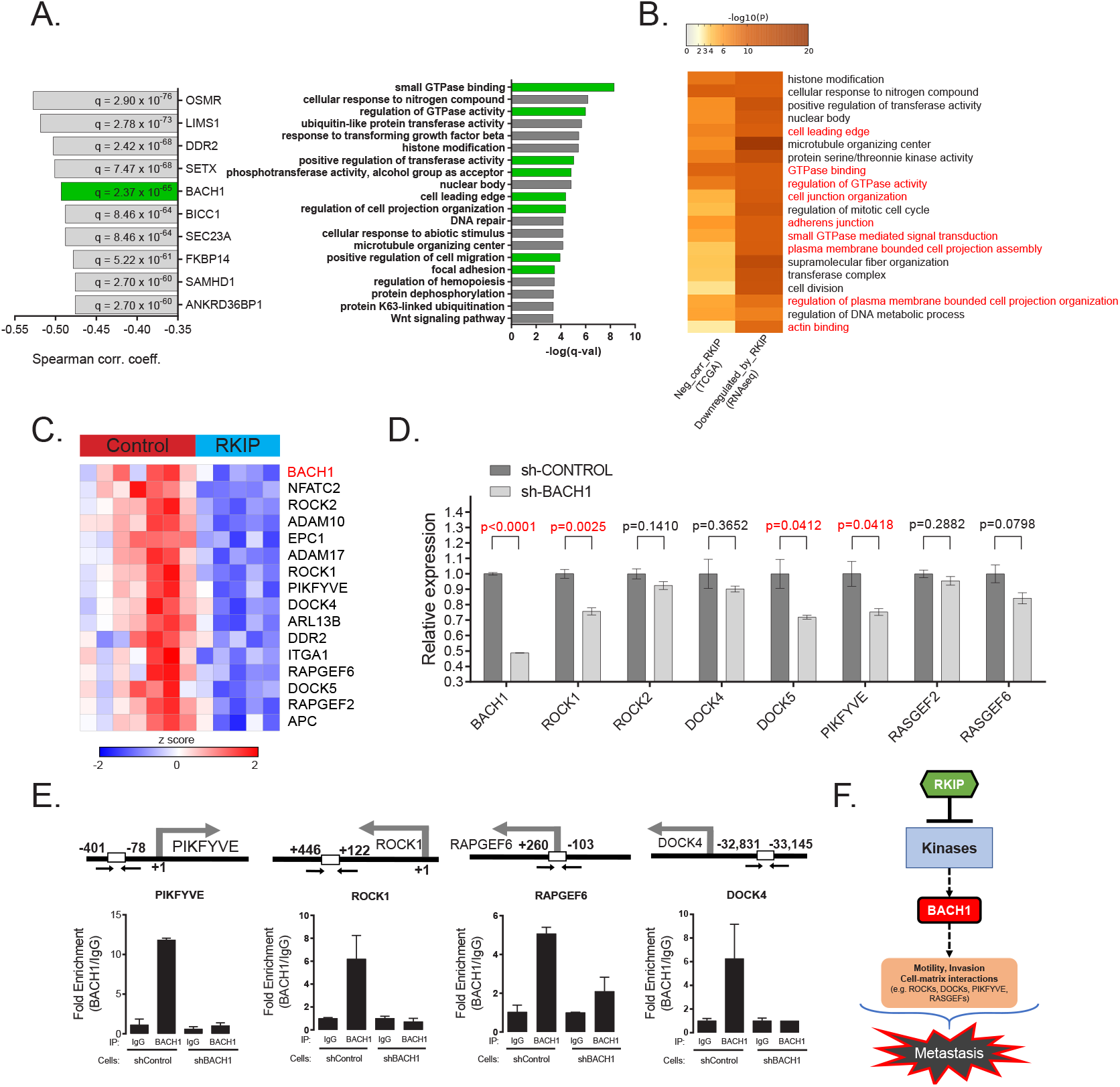
RKIP regulates a clinically relevant set of motility-related genes driven by the pro-metastatic transcription factor BACH1. **A**, Left panel: Top 10 genes negatively correlated with RKIP expression in TCGA BRCA samples (provisional, n=1100), ranked by Spearman correlation coefficient. Right panel: Gene sets enriched in genes negatively correlated with RKIP in TCGA BRCA set. **B**, Gene sets commonly enriched in genes negatively correlated with RKIP in TCGA BRCA set and genes downregulated by RKIP in the RNA-seq study. **C**, A set of differentially expressed motility genes and BACH1 ex-pression in control (n=7) vs. RKIP-expressing (n=5) BM1 tumors. **D**, qRT-PCR analysis of control (n=3) and shBACH1-ex-pressing (n=3) BM1 tumors, demonstrating downregulation of motility gene expression when BACH1 levels are reduced. Student’s t-test, two-tailed. **E**, Chromatin immuno-precipitation analysis of BACH1 binding in the promoter regions of the motility genes in BM1 cells. Mean ± s.e.m of two independent experiments. **F**, Summary diagram showing regulation of BACH1 and motility gene expression by RKIP.

The RNA-seq analysis also revealed upregulation of genes related to mitochondrial metabolism and oxidative phosphorylation in RKIP-expressing BM1 tumors (Supplementary Fig. 1C). Moreover, the mitochondrial gene targets of RKIP positively correlated with RKIP, while negatively correlating with BACH1 expression in breast cancer patients (Supplementary Figs. 1E and 2A). Our previous work similarly showed that reducing BACH1 expression in TNBC increased oxidative phosphorylation in mitochondria, mirroring the RKIP phenotype^25^. This prompted us to investigate whether BACH1 is responsible for regulating motility-related gene targets of RKIP as well. Indeed, BACH1 expression positively correlates with RKIP-inhibited motility genes in both patient samples and xenograft TNBC tumors (Supplementary Fig. 2A). ENCODE ChIP-seq analysis shows BACH1 binding to the promoter regions of the motility genes (Supplementary Fig. 2B)^26^. To confirm that BACH1 transcriptionally regulates metastatic motility-related genes in TNBC cells, we performed qRT-PCR in human TNBC cells and tumors expressing shRNAs against BACH1 (Fig. 1D and Supplementary Fig. 2C,D) and ChIP assays to demonstrate direct BACH1 binding to motility gene promoters (Fig. 1E). These findings establish BACH1-controlled motility genes as pro-metastatic targets of RKIP and illustrate the clinical relevance of this regulatory system to TNBC patients (Fig. 1F).

### RKIP targets multiple kinases in the stress MAPK network

RKIP inhibits the activity of Raf and other kinases in cultured cells (reviewed in^27^). To identify kinases targeted by RKIP in TNBC tumors, we analyzed changes in kinase expression and activity by MIB-MS analysis^3^. To capture and quantify functional kinases in tumors, tumor lysates were exposed to kinase inhibitors covalently linked to Sepharose beads (MIBs) followed by mass spectrometry. Of the 250 captured kinases that were present in both control and RKIP-expressing BM1 tumors from mouse xenografts, RKIP significantly altered the functional capture of 30 kinases (Fig. 2A). Consistent with its role as a kinase suppressor, RKIP inhibited most of these kinases (23) including the previously identified RKIP target ERK2^28^. The kinases targeted by RKIP were distributed across multiple branches of the kinome tree and not limited to a specific kinase class (Fig. 2B).

**Fig. 2:**
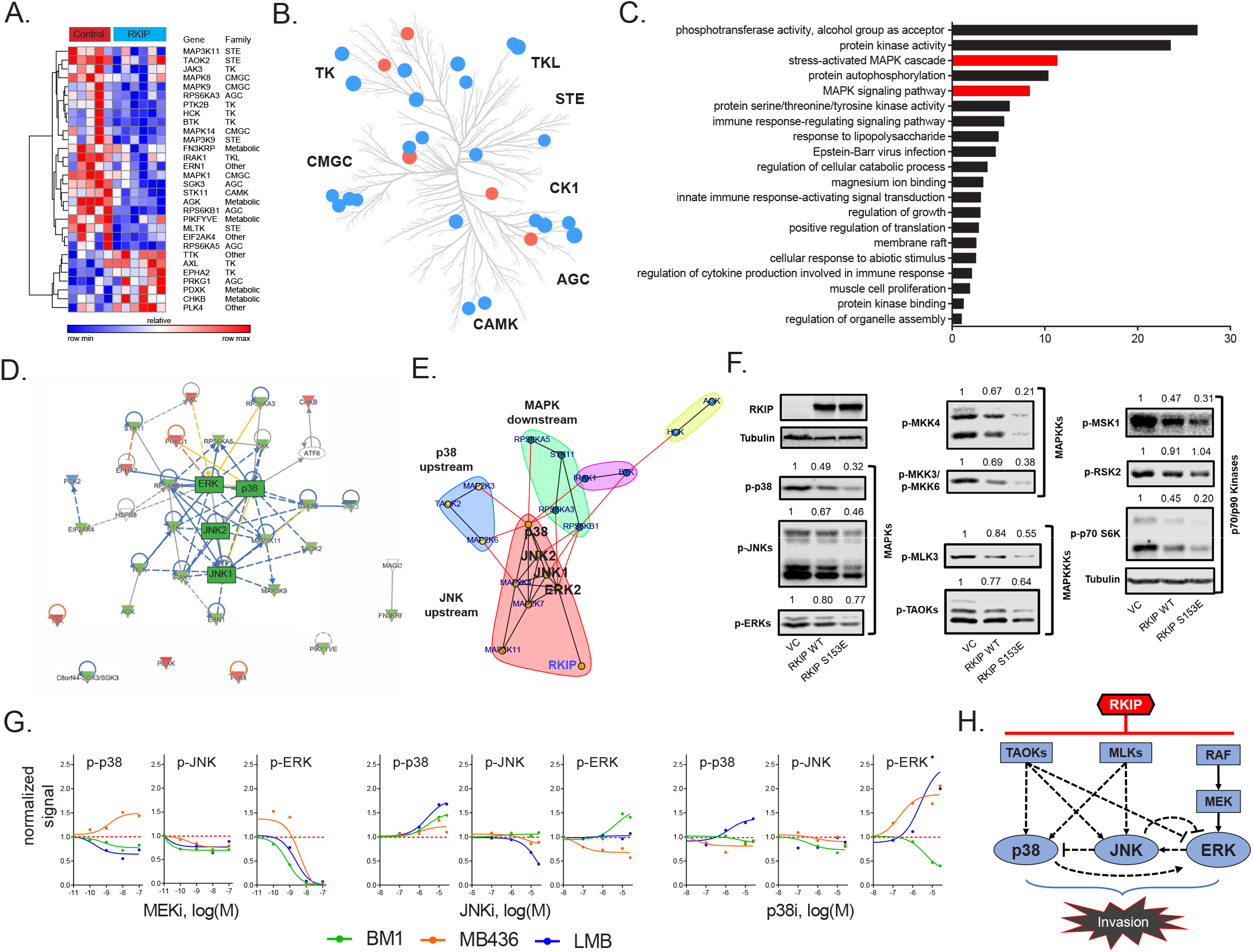
RKIP reprograms tumors by reducing signaling capacity of a network instead of targeting a single node. **A**, Multiplexed inhibitor beads - mass spectrometry (MIB-MS) analysis of n=5 control and n=6 RKIP-expressing BM1 tumors, showing 23 kinases with reduced activity and 7 kinases with enhanced activity by exogenous RKIP expression. Student’s t-test, p < 0.05. **B**, Kinome tree displaying the distribution of kinases targeted by RKIP across different families of kinases. Blue: activity reduced by RKIP (n=23), Red: activity enhanced by RKIP (n=7). **C**, Gene set enrichment analysis of the 23 negatively regulated kinases by Metascape. Stress-induced mitogen activated protein kinase (MAPK) related gene sets are highlighted in red. d, Ingenuity® Pathway Analysis (IPA) of the RKIP target kinases centered around MAPKs p38, JNK, and ERK. **E**, Direct protein-protein interaction network and community analysis showing the core of the RKIP kinase network. **F**, Downregulation of the stress MAPK network by wild type RKIP and the Raf-binding S153E mutant in BM1 cells in vitro. Quantification with respect to Tubulin signal. Representative of two independent experiments. **G**, Effect of inhibiting an individual MAPK on p38, JNK, and ERK activity across multiple cell lines under anisomycin-induced stress conditions. Each dot represents the phospho-kinase signal in a different cell line (also see Supplementary Fig. 4). **H**, Diagram summarizing the interactions within the RKIP-regulated stress MAPK network in anisomycin-induced BM1 cells. Kinase interactions are determined by using small molecule inhibitors or siRNAs against the kinases in the network. The TAOK-p38 interaction is observed in cells treated with a cocktail of siRNAs against all three TAOKs (siCombo), whereas TAOK-JNK and TAOK-ERK interactions were obserbed by siRNAs against TAOK1 and TAOK2 respectively.

Functional analysis of the downregulated kinases using Metascape^29^ showed enrichment for stress kinase signaling (Fig. 2C). Ingenuity® pathway analysis indicated that most of these kinases are functionally related, and the stress MAP kinases (JNK, p38) as well as ERK comprise the core of the network (Fig. 2D). Community analysis identified three main protein-protein interaction subnetworks within the extended MAPK family including kinases upstream of p38 (TAOK2, MKK3, MKK6), kinases upstream of JNK (MLK1, MLK3, MKK4); and p70/p90 kinases downstream of MAPKs (MSK1, RSK2, and p70/85 S6K1) (Fig. 2E). We validated inhibition of stress-activated MAPKs by exogenously expressed RKIP or a constitutively active mutant S153E RKIP^9^ in human TNBC cell lines (BM1, MDA-MB-436) stimulated with a stress signal, the protein synthesis inhibitor anisomycin (Fig. 2F and Supplementary Fig. 3A). We confirmed the importance of stress MAPK signaling for invasion^30^ using small molecule inhibitors of p38 and JNK (Supplementary Fig. 3B). These results indicate that the three MAPKs (JNK, p38 and ERK) as well as their upstream regulators and downstream effectors comprise the core RKIP-regulated network that drives metastasis in TNBCs.

To understand how RKIP targets the MAPK signaling network, we identified upstream regulators of MAPKs inhibited by RKIP in tumors. MIB data from BM1 xenografts revealed three members of the MAP3K family reported to activate both p38 and JNK signaling: TAOK2, MLK1 and MLK3^30–32^ (Fig. 2A,F and Supplementary Fig. 3A). Treatment of TNBC cells with selective inhibitors or siRNAs against these MAPKKKs confirmed their ability to inhibit downstream MAPKs. Specifically, TAOKs primarily regulate p38 but may also activate JNK in TNBC cells, and MLK1,3 primarily activate JNK in TNBC cells but can also activate p38 dependent on the stimulus and cell type (Supplementary Fig. 4). As expected, TNBC invasion was blocked by inhibition of either MLKs or TAOKs (Supplementary Fig. 3C,D) (see also^33^). These data suggest that RKIP inhibits the upstream TAOKs and MLKs in addition to Raf in TNBC tumors, thereby preventing activation of the pro-invasive stress MAPKs p38 and JNK in TNBC cells.

To determine whether targeting a single MAPK is sufficient to mimic RKIP and attenuate signaling by the other MAPKs, we inhibited p38, JNK, or MEK individually. A dose response for MEK inhibition under stress (anisomycin) conditions revealed robust suppression of pERK but limited inhibition of JNK or p38, respectively, in BM1 cells, and a comparable pattern was observed with other cell lines (Fig. 2G and Supplementary Fig. 4). Similar restricted inhibition profiles were observed when we used p38 or JNK inhibitors alone (Fig. 2G). This analysis suggests that targeting any one of the three MAPKs alone is insufficient to mimic RKIP and reduce signaling from all three MAPKs. Our results indicate that, in contrast to the common therapeutic practice of fully inhibiting individual kinases at the maximum-tolerated dose, RKIP functions like a low dose, non-toxic drug combination that reduces the activity of several key kinases within a driver network to inhibit metastasis (Fig. 2H).

### A low dose four-drug combination reduces MAPK network signaling capacity and suppresses tumor cell invasion without altering growth

Three aspects of RKIP regulation provide strategic guidance for anti-metastatic therapy. First, as noted above, RKIP suppresses the signaling capacity of multiple kinases within a key driver network, the stress MAPK network. Second, the kinases that are linked to metastasis ranged from high to low functional capture, indicating that the degree of kinase activity does not correlate with metastatic potential (Fig. 3A). Finally, the effective decrease in kinase capture for RKIP targets was generally less than 30% (Fig. 3B). We then determined whether we could mimic the action of RKIP using drugs to inhibit MAPK network signaling.

**Fig. 3:**
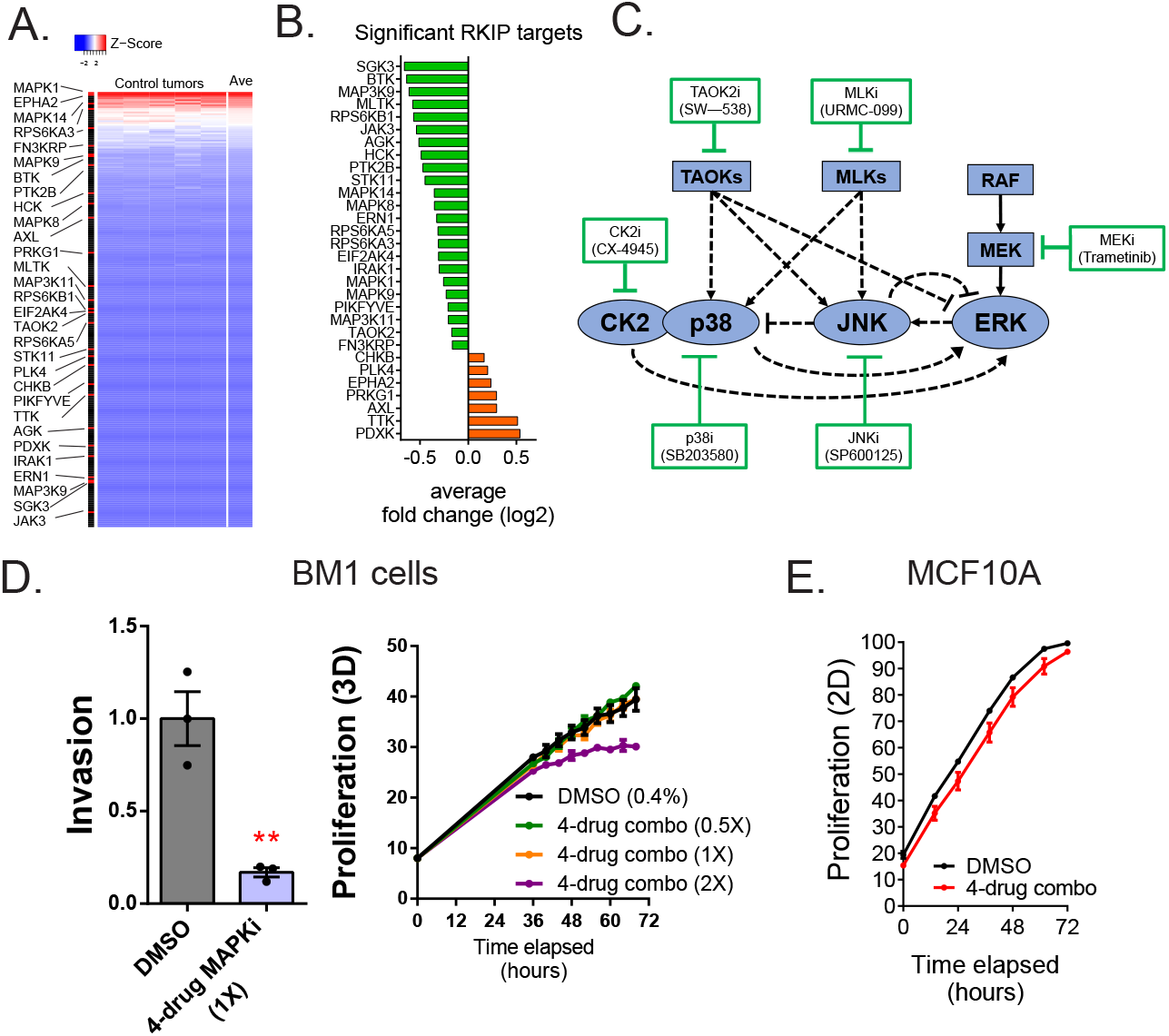
A low dose four-drug combination reduces MAPK network signaling capacity and suppresses tumor cell invasion without altering growth. **A**, Heatmap demonstrating the distribution of kinases targeted by RKIP in an activity ranked list of all 250 kinases cap-tured. Top-to-bottom represents high-to-low ranking of MIB-captured kinases in the control cases (n=7). **B**, Average effect size of changes in kinase activity induced by RKIP in RKIP-expressing BM1 tumors vs. control. **C**, Diagram of the network used for modeling and the small molecule inhibitors used in the high-throughput invasion assays for potential drug combinations. **D**, Chemotactic invasion assay (left panel) and 3D proliferation assay (right panel) showing that the 4-drug MAPK inhibitor combination (4D-MAPKi) blocks invasion of BM1 cells without inhibiting their growth. Mean ± s.e.m of n=3 technical replicates per experimental group. Two-tailed student’s t-test for the invasion assay. Two-way ANOVA test with Tukey’s multiple comparison correction for the proliferation assay. **E**, 4D-MAPKi is not toxic to immortalized human mammary epithelial MCF10A cells. *p<0.05, **p<0.01, ***p<0.001, ****p<0.0001.

We sought a drug combination that, like RKIP, reduces invasion but not cell growth. We initially assessed combinations of 6 kinase inhibitors that target different nodes in the MAPK signaling network using BM1 cells (Fig. 3C). In addition to the MEK, JNK, p38 and MLK inhibitors used above, we also tested CX-4945 (Silmitasertib), an inhibitor of Casein Kinase 2 (CK2) which is involved in p38 and ERK signaling^34–36^, as well as SW-538, a more broad-based inhibitor that blocks some kinases that act on the MAPK network such as TAOK2, Raf1, JNK1, HGK, and GSK3b^37^.

We monitored invasion as well as growth of cancer cells in 3D culture using a high-throughput invasion assay of nuclear-labeled BM1 cells (BM1-NucLight Red). For four drugs (SB203580, SP600125, Trametinib, URMC-099), there was a minimal dose at which the proliferation rates were unaffected while the invasive capabilities of the cells were at least partially blocked (Supplementary Fig. 5A). We then tested these drugs at these minimal dosages for their combinatorial effect on invasion. Out of all the dual combinations tested, only two demonstrated a combinatorial effect on invasion without a significant effect on proliferation: p38i+JNKi and MEKi +MLKi (Supplementary Fig. 5B). Addition of a third inhibitor to these dual combinations did not improve the anti-invasive efficacy, demonstrating that the combined effect of multiple MAPK inhibitors is not necessarily additive (Supplementary Fig. 5C).

A four-drug combination consisting of p38i, JNKi, MEKi, and MLKi was more successful than either of the dual combinations at inhibiting both MAPK signaling (JNK, MEK, p38) and cell invasion across multiple cell lines and stimuli, but had no effect on proliferation (Fig. 3D and Supplementary Fig. 5D-G). Notably, 4D-MAPKi was not toxic to normal human mammary epithelial cell lines MCF10A and 184A1 (Fig. 3E and Supplementary Fig. 5H). These findings suggest that the 4D-MAPKi combination is an effective, well-tolerated anti-invasive therapy that mimics the strategy by which RKIP inhibits the MAPK network.

### The four-drug combination suppresses metastasis, inhibits expression of pro-metastatic motility genes, and increases survival

We then determined whether 4D-MAPKi blocks metastasis of TNBC tumors *in vivo*. Using mouse LMB cells in a syngeneic TNBC model, we performed dose-response studies with individual drugs to determine the highest dose at which growth of the primary tumor is unaffected (Supplementary Fig. 6A). Based on this analysis, we chose a 4D-MAPKi combination of 10 mg/kg for p38i, JNKi, MLKi, and 0.5 mg/kg for MEKi (1X) for *in vivo* studies.

At the 1X dose, the 4D-MAPKi combination significantly inhibited primary tumor growth in both syngeneic LMB tumors (Fig. 4A) and xenograft BM1 tumors (Fig. 4B) in a dose-dependent manner without obvious toxicity as all mice retained the same body weight (Supplementary Fig. 6B). In order to mitigate the confounding effect of primary tumor growth on metastasis and maximize metastatic burden, we employed tail-vein or intracardiac injection models of experimental metastasis. Both undiluted (1X) and 50% diluted (0.5X) 4D-MAPKi suppressed metastatic lung colonization in syngeneic LMB tumors in a dose-dependent manner (Fig. 4C,D). Treatment of mice with 4D-MAPKi for only 2 days following tumor cell injection still caused significant reduction in the overall metastatic burden ~5 weeks later, suggesting that the inhibitor combination suppresses early steps of metastasis related to invasion and extravasation (Fig. 4E). Human BM1 tumors responded better to the 4D-MAPKi treatment than LMB tumors, as even the half dose (0.5X) potently inhibited bone metastasis (Fig. 4F,G). The 4D-MAPKi combination also improved survival of metastatic BM1-bearing mice following a three-week treatment (Fig. 4H). 4D-MAPKi, like RKIP, blocked induction of a significant fraction of both BACH1 and motility genes in syngeneic LMB tumors (Fig. 4I). These findings indicate that the low dose drug combination, 4D-MAPKi, suppresses TNBC metastasis by reducing the signaling capacity of the stress MAPK network that transcriptionally activates metastatic genes, thereby increasing survival.

**Fig. 4:**
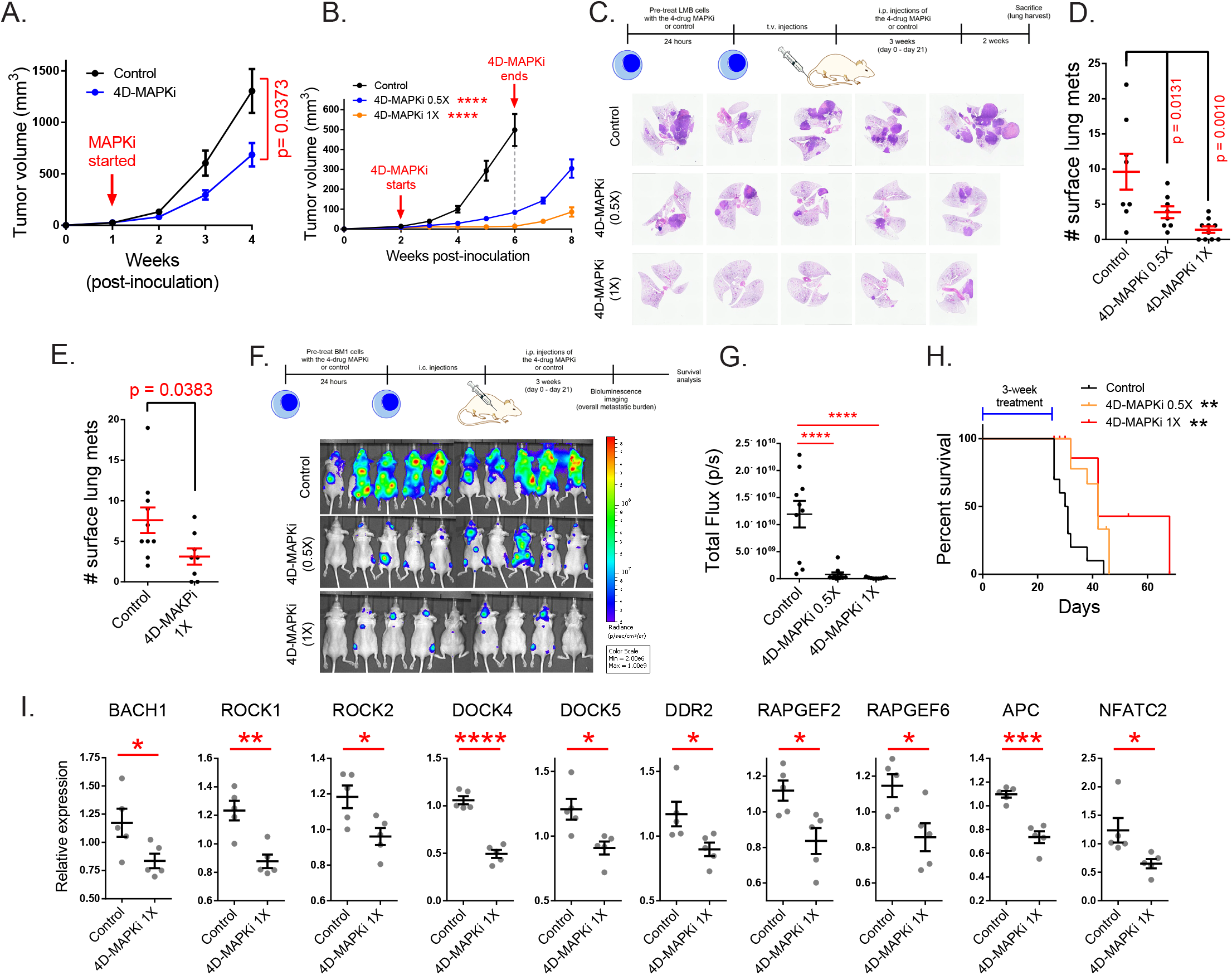
The four-drug combination suppresses metastasis, increases survival and inhibits expression of pro-metastatic motility genes. **A**, Effect of MAPKi treatment on the primary LMB tumor growth. Mean ± s.e.m. of n=5 biological replicates per experimental group. Two-way ANOVA test. **B**, Primary BM1 tumor growth in mice treated with 4D-MAPKi combination for 4 weeks. Mean ± s.e.m of n=8 control tumors, n=8 4D-MAPKi(0.5X) treated tumors, and n=6 4D-MAPKi(1X) treated tumors. Two-way ANOVA test with multiple comparisons at week 6. **C**, 4D-MAPKi combination reduces metastatic tumor burden in the lungs of LMB syngeneic mouse model of TNBC. H&E staining demonstrates the metastatic lesions in cross-sections of the lungs in mice treated with 1X (undiluted, n=10 biological replicates) 4D-MAPKi, 0.5X (diluted, n=8) 4D-MAPKi, or the control (vehicle, n=8). **D**, Quantification of the visible metastatic lesions on the lung surface. Mean ± s.e.m, one-way ANOVA test with Dunnett’s correction for multiple testing. **E**, Tumor burden in the lungs of LMB syngeneic mice after 2 days (2 doses over 48 hours, on day 0 and day 1) of 4D-MAPKi treatment. Mean ± s.e.m. of n=10 control tumors and n=8 MAPKi(1X) treated tumors. Unpaired two-tailed student’s t-test. **F**, BM1 metastatic tumor burden in the bones of athymic nude mice treated with 4D-MAPKi at 0.5X and 1X, or control. **G**, Quantification of the metastatic burden in (E). Mean ±ns.e.m. of n=10 control, n=10 diluted 0.5X 4D-MAPKi, and n=9 undiluted 1X 4D-MAPKi.One-way ANOVA test with Dunnett’s correction for multiple testing. **H**, Overall survival of xenograft mice injected with BM1 cells via the intracardiac route after 3 weeks of 4D-MAPKi treatment. Log-rank (Mantel-Cox) test. **I**, Expression of BACH1 and motility genes in LMB tumors treated with 4D-MAPKi. Mean ± s.e.m. of n=5 biological replicates per experimental group. Two-tailed student’s t-test. *p<0.05, **p<0.01, ***p<0.001, ****p<0.0001.

### Multi-drug combination inhibits different MAPK network topologies

To understand why the 4-drug combination is so effective, we performed further experiments as well as phenomenological (observation-based) modeling of the MAPK network. Characterization of the cross-talk between different MAPKs in the RKIP-regulated network revealed that each of three human or mouse TNBC cell lines has a unique MAPK network topology (compare Fig. 2H to Supplementary Fig.7A; also see Supplementary Figs. 4A-C). Here we define topology as interactions between different nodes within the network that could be positive, negative, direct, or indirect. Although these network analyses are not exhaustive, our results illustrate differences in the cellular network topologies. To determine whether environmental stimulus also makes a difference in MAPK network topology, we treated BM1 cells with either anisomycin or serum, and then probed cellular response with the p38 inhibitor (p38i). As noted above, inhibition of p38 activation by p38i did not significantly inhibit JNK in anisomycin-treated cells, whereas it actually activated JNK under serum conditions (Fig. 2H and Supplementary Fig. 4D). These results suggest that MAPK network topology can differ between cell lines, or even within in the same cell line based on stimulus, and justify the importance of considering network topology in determining treatment efficacy.

In order to understand the effect of low dose, multi-drug treatment of metastatic cancer, we constructed a simple steady state model of the core MAPK network. Since a description based on dynamical systems theory is beyond the scope of the present work, we utilized a network flow approach^38^. Our goal was to determine the effect of introducing crosstalk into a system where a stress signal flows through three spigots (TAOK/p38, MLK/JNK and RAF/ERK pathways) and funnels into a combined output measured as BACH1 transcription. The maximal flow through the network is defined by the condition when no drugs have been administered. A node represents a kinase whose activity determines its capacity to absorb the inflow signal, and this capacity can be reduced by drug treatment. In this analysis, we assume the inflow signal equals the outflow signal.

We utilized the anisomycin-stimulated MAPK network in BM1 cells as the basis for our model (Fig. 5A, Network 1). If we modeled the network from other cell lines, the detailed topology would be different but the same general principles apply. We illustrate this point by generating another related MAPK network based on Network 1 but lacking the positive crosstalk (Fig. 5B, Network 2). Comparison of treatment with single MAPK inhibitors (p38i, JNKi or MEKi) to 4D-MAPKi (dose restricted to ≤30% inhibition for each kinase targeted) reveals at least 60% BACH1 suppression for 4D-MAPKi and p38i in Network 1; by contrast, BACH1 levels following JNKi or MEKi treatment were only minimally reduced (Fig. 5C). A similar result is obtained for Network 2, although in this case JNKi has no significant effect on BACH1 levels (Fig. 5D). These results from our models suggest that, while response to individual inhibitors may vary because of differences in network topology, 4D-MAPKi is more robust in inhibiting network output across different topologies.

**Fig. 5:**
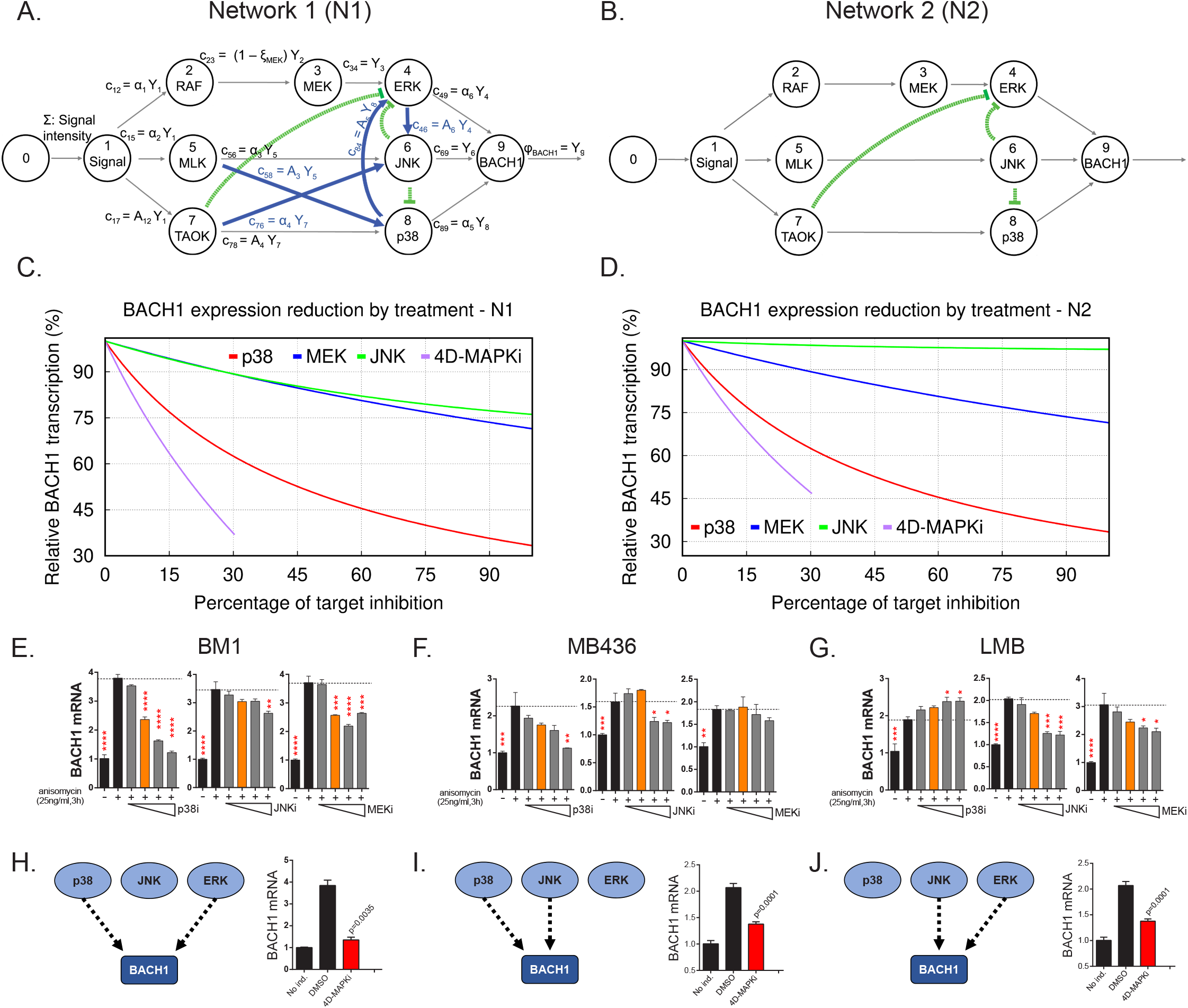
Multi-drug combination inhibits different MAPK network topologies. **A**, The topology of the stress response of the core MAPK driver network activating BACH1 transcription, as output, in BM1 cells. This network topology, termed N1, is composed of multiple kinase signaling pathways responsible for activating BACH1. The nodes of the signaling network, represented by circles, are kinases within the network. The arrows directed toward nodes indicate the inflow from a signal or an active kinase at the upstream node. The product of a node, resulting from the interaction between the upstream signal and downstream kinases, is denoted by arrows leaving the node. The pathways are indicated by black arrows while the crosstalk between different pathways is denoted by blue and green arrows. The non-linear repression of one node by another is represented by the green lines with bars directed toward the repressed component. **B**, A hypothetical BACH1 stress response driver network, denoted as N2, that has no crosstalk between its individual pathways. The interpretation of its symbols is the same as in Fig. 5A. **C**, Graph depicting predicted downregulation of BACH1 following cell treatment with specific inhibitors relative to maximal stress-induced BACH1 transcription in cells with N1. The y axis shows the percentage of maximal BACH1 transcription, and the x axis denotes the percent inhibition of each kinase targeted by a drug or drug combo relative to the maximal inhibitor dose (set at 100% inhibition). Relative BACH1 transcription in response to p38i, MEKi, JNKi, or the 4D-MAPKi drug combo is indicated in red, blue, green, or purple. **D**, Graph depicting predicted downregulation of BACH1 following cell treatment with specific inhibitors relative to maximal stress-induced BACH1 transcription in cells with N2. The y axis shows the percentage of maximal BACH1 transcription, and the x axis denotes the percent inhibition of each kinase targeted by a drug or drug combo relative to the maximal inhibitor dose (set at 100% inhibition). Relative BACH1 transcription in response to p38i, MEKi, JNKi, or the 4D-MAPKi drug combo is indicated in red, blue, green, or purple. **E-G**, Single agent dose-response experiments demonstrating that BACH1 expression is activated by different MAPKs in different cell lines. Orange bars indicate the final dosage of an individual inhibitor used in 4D-MAPKi. Mean ± s.e.m of n=3 technical replicates. One-way ANOVA with multiple testing correction. **H-J**, The network-targeting 4D-MAPKi is able to decrease BACH1 expression across all three cell lines even though BACH1 is regulated by a different set of MAPKs in each cell line. (Left panels) Diagrams summarizing BACH1 regulation by MAPKs in each TNBC cell line. (Right panels) 4D-MAPKi blocks BACH1 mRNA expression in anisomycin-induced cells. The bar-graphs are representative of two independent experiments in each cell line. Mean ± s.e.m of n=3 technical replicates. *p<0.05, **p<0.01, ***p<0.001, ****p<0.0001.

To test this prediction, we assessed BACH1 gene expression as a measure of network output in the three TNBC cell types with different network topologies. Dose-response studies with individual MAPK inhibitors showed that BACH1 expression can be regulated by either ERK, JNK or p38 in at least one TNBC cell line (Fig. 5E-G). However, as our model predicted, no single MAPK inhibitor even at maximum dose effectively reduced BACH1 expression across the three cell lines (Fig. 5H-J, left panels). By contrast, the low dose 4D-MAPKi combination attenuated both BACH1 gene or protein expression in all human and mouse cell lines tested (Fig. 5H-J, right panels and Supplementary Fig. 7B). These data indicate that 4D-MAPKi more effectively regulates BACH1 expression across cells with different MAPK network topologies than single high-dose inhibitors.

### Limiting the extent of kinase inhibition at multiple nodes reduces network output and prevents compensatory network activation

Maximum tolerated dose regimens can yield promising responses in certain patients.^2–4^ In terms of our mathematical model, when all pathways are independent and operating at their maximal capacity, reducing or eliminating any one of them restricts the output flow. In this scenario, a single target therapy can potentially be effective at reducing output. However, if flow from the single node is eliminated, then excess flow from the initial functional network could end up activating a different functional network, leading to a compensatory increase in overall output^39^ (see CKN, compensatory kinase network in Fig. 6A).

**Fig. 6:**
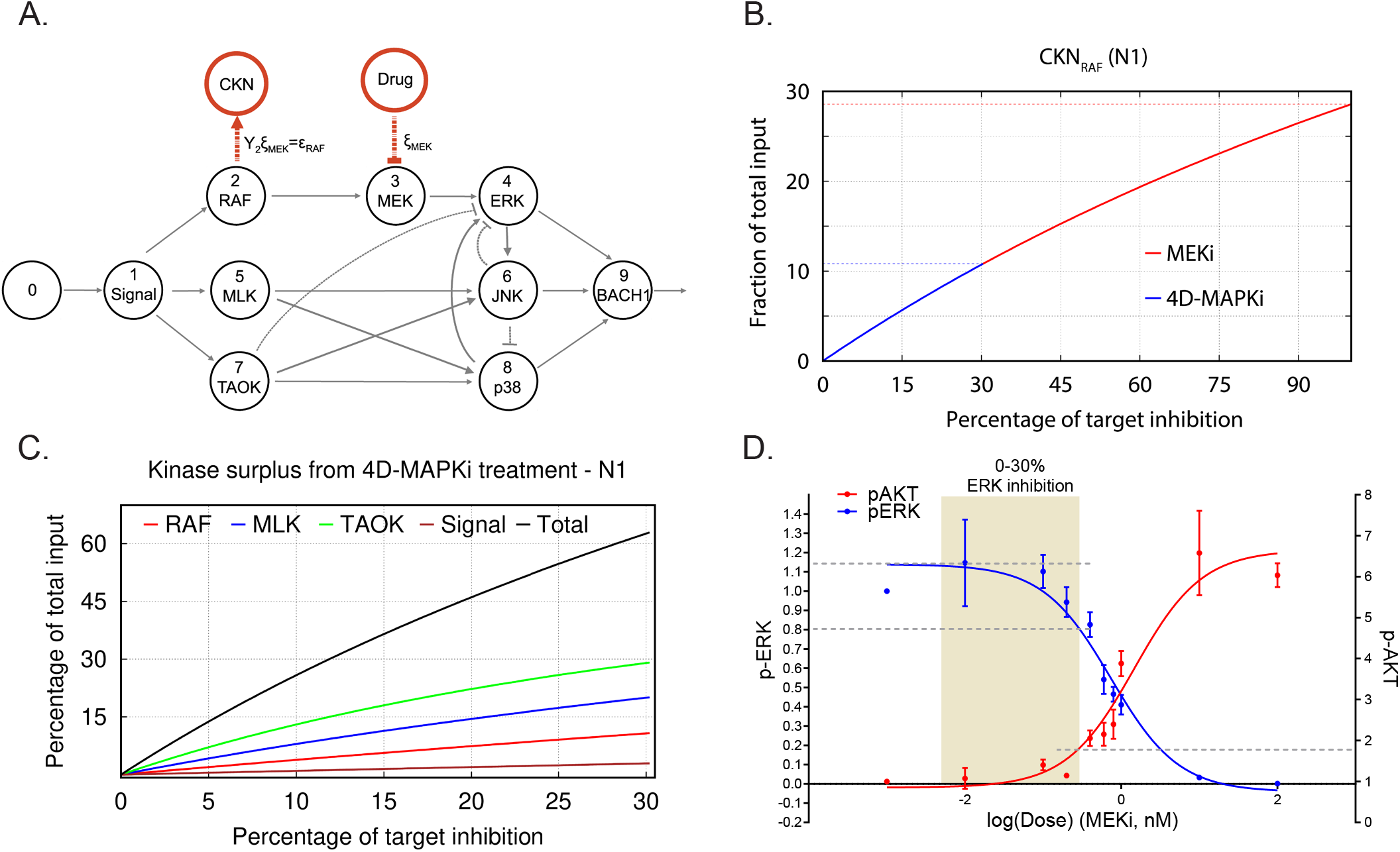
Limiting kinase inhibition at multiple nodes reduces network output and prevents compensatory network activation. **A**, N1 illustrating the case of a treatment (MEKi) targeting node 3 (MEK) and activation of a compensatory kinase network (CKN) linked to upstream node 2 (RAF) (red circle). A comparable diagram can be generated for each upstream node to describe surplus signal activating a distinctive CKN. The percentage of reduction on the activity at a target node because of a treatment dose *x* is indicated by ξ. This inhibition causes reduction in the flow of product to node 3, denoted by c_23_=(1−ξ_MEK_)Y_2_ (see Fig. 6A). **B**, Graph depicting surplus signal from RAF as a fraction of the total input signal Σ following treatment by MEKi or 4D-MAPKi of cells with N1. The x axis denotes the percent inhibition of each kinase targeted by inhibitors relative to the maximal inhibitor dose (set at 100% inhibition). The y axis is the fraction of surplus RAF signal generated following drug treatment of cells relative to total input signal. MEKi, red; 4D-MAPKi, blue. **C**, Graph depicting surplus kinase signal as a fraction of the total input Σ following treatment by 4D-MAPKi of cells with N1. The surplus is a consequence of the congestion of each direct pathway causing an insufficient absorption of the stress input by the driver network and its redirection towards a compensatory network. The x axis denotes the percent inhibition of each kinase targeted by 4D-MAPKi relative to the maximal inhibitor dose (set at 100% inhibition). The y axis is the fraction of surplus kinase signal generated following 4D-MAPKi treatment of cells relative to total input signal. RAF (surplus from MEKi), red; MLK (surplus from JNKi), blue; TAOK (surplus from p38i), green; and Signal (surplus from MLKi), brown; Total (sum of all surplus signals), black. **D**, Dose-response curves showing activation of the compensatory PI3K network, monitored by p-AKT levels, when EGF-induced BM1 cells are treated with increasing doses of MEKi. Mean ± s.e.m. of n=3 independent experiments.

To address the role of surplus signal flow following inhibitor treatment, we determined whether different degrees of inhibition of MEK would yield different outcomes with respect to activation of compensatory driver networks. As a model system, we analyzed induction of an epidermal growth factor (EGF) receptor-PI3K feedback loop in response to MEK inhibition of EGF-stimulated BM1 cells. Activation of this compensatory PI3K pathway was previously reported to restrict the efficacy of MEK inhibitors in basal subtypes of breast cancer^40^. Following maximal inhibition of MEK in our model, the surplus flow from Raf approaches 30% of the initial input signal that can be funneled to a compensatory kinase network such as PI3K/Akt (Fig. 6B). When we modeled 4D-MAPKi, on the other hand, it only diverted ~10% of the incoming stress signal to a compensatory kinase network (CKN) because we restricted MEK/ERK inhibition to less than 30%. This type of analysis, carried out for each of the kinases in the MAPK network, shows that the combined surplus of kinase signal following 4D-MAPKi inhibition approaches 60% (Fig. 6C). However, this signal is dissipated among multiple kinases such that the resulting surplus from each kinase is insufficient to activate compensatory networks.

To test these predictions experimentally, we generated dose response curves to assess the relationship between degree of MEK inhibition (assessed by pERK) and induction of PI3K (assessed by pAkt). The results show that Akt activation has a threshold effect with minimal change in activity until ERK is inhibited by ~30% (Fig. 6D). Consistent with this finding, when we tested 4D-MAPKi in EGF-stimulated cells, ERK signaling was robustly inhibited (~75%) and Akt was maximally activated (Supplementary Fig. 7C). By contrast, in serum-stimulated cells, 4D-MAPKi inhibited ERK by only ~40% and Akt activity was actually reduced (Supplementary Fig. 7D). Taken together, these results show that limiting ERK inhibition, as RKIP does, is an effective mechanism to avoid compensatory Akt activation above background.

Taken together, these experimental and mathematical analyses suggest that effective inhibition can be accomplished by targeting several nodes belonging to different pathways within the same driver network. This will reduce the flow through multiple pathways of the network and its resulting output, decreasing the efficacy of the driver network. Reduction of overall output, however, creates a surplus signal that cannot be accommodated by other kinases within the network and, instead, is directed towards compensatory driver networks. Thus, it is important to keep the overall surplus dissipated among multiple nodes, and the surplus signal from each kinase below the threshold for activation of its compensatory network. Of note, since our goal is to suppress metastasis but not growth, this partial inhibition which leaves the growth network intact is still effective. Together, these studies suggest that 1) multi-kinase targeting is more effective than single kinase inhibition across different cells and environmental stimuli; and 2) low inhibitor doses are less likely than high inhibitor doses to trigger feedback activation of compensatory networks.

### The BACH1/motility gene axis, targeted by RKIP and 4D-MAPKi, is associated with multiple cancers and metastasis suppressors

To understand the clinical significance of the MAPK network suppressor (RKIP) and the MAPK network output (BACH1 and motility-related genes), we looked at their relative expression in the TCGA database. Remarkably, stratifying breast cancer patients by high RKIP and low BACH1 expression or vice versa reveals a striking inverse association of RKIP with BACH1 and the motility-related genes in ~60% of patients (Fig. 7A). These data suggest that the RKIP/BACH1/motility gene axis identifies breast cancer patients who would be therapeutic candidates for 4D-MAPKi treatment. Enrichment of the motility-related target genes identified in the present study extended beyond breast cancer. The same gene families were also inversely correlated with RKIP in other solid TCGA cancer types (Fig. 7B,C) including pancreatic, ovarian, lung, head and neck, and colorectal. Of note, in cancers where no correlation to these specific motility-related genes was observed, we noticed a strong correlation to other members of the same gene families (Fig. 7C). Finally, expression of other experimentally validated metastasis suppressors (BRMS1, ARGHDIA, NME1, and DRG1)^8^ also negatively correlated with motility-related gene sets (Fig. 7D,E). These clinical analyses suggest that BACH1-regulated motility-related machinery is a hallmark of metastasis that is targeted by multiple physiological suppressors such as RKIP.

**Fig. 7:**
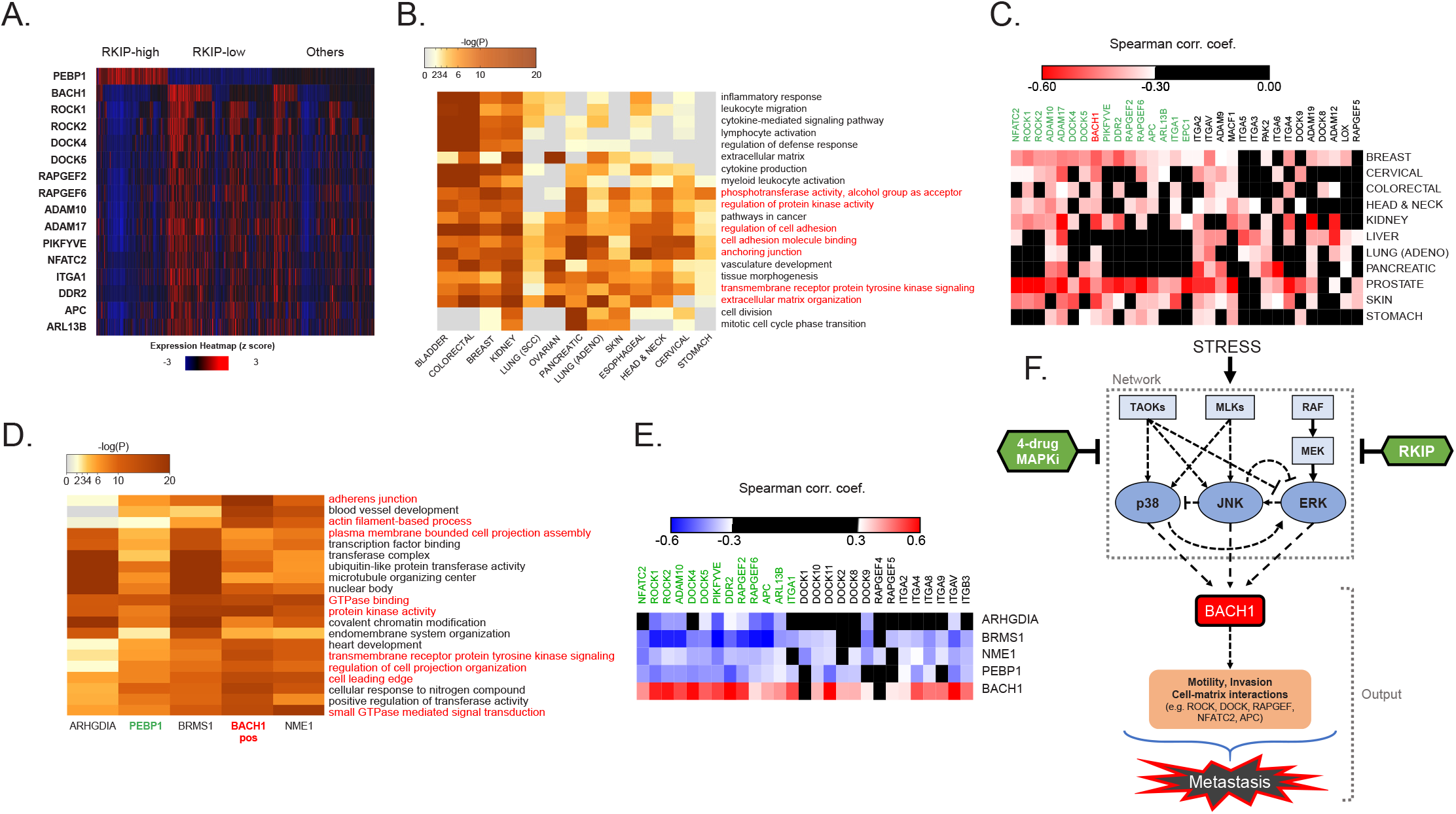
The BACH1/motility gene axis, targeted by RKIP and 4D-MAPKi, is associated with multiple cancers and metastasis suppressors. **A**, Expression of RKIP (PEBP1), BACH1, and the downstream motility genes in each TCGA BRCA patient (n=1100), grouped by RKIP status. RKIP-high: z-score > 0.5 (n=274), RKIP-low: z-score < −0.5 (n= 414), Others: −0.5 < z-score < 0.5 (n=412). **B**, Gene sets enriched with genes negatively correlated with RKIP across multiple TCGA cancer types. **C**, Spearman correlation coefficients for BACH1 or motility genes relative to RKIP in TCGA cancers. Coefficient cutoff of −0.3. **D**, Motility-related gene sets enriched for genes that negatively correlate with the indicated metastasis suppressors, but positively correlate with BACH1 in the TCGA BRCA set. **E**, Spearman correlation coefficients for BACH1 or motility genes relative to RKIP (PEBP1) or other metastasis suppressors in TCGA BRCA. Coefficient cutoff of −0.3. **F**, Diagram summarizing the stress MAPK kinase N1 network that regulates metastasis in breast cancer. Stress activates a network of MAPKs that interact via crosstalk. RKIP and the RKIP-mimicking drug combo 4D-MAPKi reduce the signaling capacity of the entire network by targeting multiple nodes. This allows for effective reduction of the metastatic output of the network, measured by the expression of pro-metastatic BACH1 and its target motility genes.

Taken together, our results show that stress as an input activates the core MAPK network, leading to induction of BACH1 expression as an output which in turn activates motility-related genes required for invasion and metastasis (Fig. 7F). Inhibition of this network by metastasis suppressors such as RKIP or a low dose 4-drug combination effectively restricts expression of invasive genes and reduces the metastatic phenotype.

## DISCUSSION

Here we propose a new approach for therapeutic targeting of metastatic disease based upon the action of physiological metastasis suppressors such as RKIP. In this study we focus on the underlying general principles of suppressor action that enable stable metastatic inhibition without triggering activation of compensatory driver networks. Instead of completely inhibiting specific nodes, suppressors reduce the signaling capacity of a driver network by partially targeting multiple kinases within the network, thereby restricting the output that promotes invasion and metastasis. We experimentally validate this approach using a 4-drug combination that acts on the core MAPK network to inhibit metastasis and promote survival in mouse TNBC models. This approach even works for different driver network topologies associated with diverse cells within tumors, conferring a higher degree of robustness to our therapeutic strategy.

Mathematical modeling using a simple steady state model illustrates the underlying concept. The model suggests that multidrug combinations are more effective at suppressing signaling across cells with different MAPK network topologies. In addition, minimizing the inhibition of each kinase enables dissipation of excess signal so that compensatory networks are not activated. Indeed, our experimental findings support these general principles, demonstrating that different TNBC cells have different MAPK network topologies and respond in diverse ways to MAPK inhibitors. Our data demonstrate that activation of pAKT after treatment of cells with MEK inhibitor is a threshold effect, further supporting the relevance of our proposal and opening a new avenue of investigation aimed at finding the range of topologies within a single driver network and their relationship to multiple compensatory driver networks.

Using databases such as TCGA combined with preclinical studies enables identification of metastasis suppressor-regulated genes that can be used as biomarkers both for development of drug combinations and identification of patients who would benefit from those therapies. Similar correlations between expression of other metastasis suppressors and motility-related genes in other tumor types suggest that this approach can have wide application to pharmaceutical treatment of metastatic disease states in multiple tissues. In particular, our findings suggest that 4D-MAPKi would be an effective treatment for metastatic breast tumors that express the motility-related genes but lack suppressors that inhibit them. We anticipate that, in addition to identifying 4D-MAPKi, this approach can be utilized to discover other multi-drug combinations that are effective.

4D-MAPKi, which targets MEK, p38, JNK, and MLK, is a novel and clinically feasible multi-drug combination^41–44^. Drugs such as these, when used as single agents, have not given durable responses thereby necessitating multi-drug approaches (e.g.^45^). Data already available from MEK inhibitor Phase 1 trials^46^ should enable an estimate of drug doses that would reduce MEK activity by less than 30%, and similar analyses can be carried out for the other kinase inhibitors. Effective treatment based on inducing functional RKIP in tumors has not been possible since RKIP regulation is complex and occurs at multiple levels including transcriptional, translational and post-translational (Yesilkanal and Rosner, 2018). By significantly expanding our knowledge of RKIP function as a tumor metastasis suppressor, this study identifies additional therapeutic targets. MLKs and TAOKs, as regulators of p38 and JNK, have not previously been implicated as a part of the RKIP-network. BACH1, which we recently identified as an inhibitor of mitochondrial metabolism that can be targeted independently in ~30% of breast tumors^25^, is shown here to be a transcriptionally-regulated and pro-invasive mediator of the stress MAPK network.

While the utility of multi-drug combinations for therapeutic treatment is widely acknowledged, the strategy proposed here differs significantly in several respects. First, we focus on reducing signaling capacity in one driver network rather than fully inhibiting single nodes belonging to one pathway. Second, our findings argue that targeting multiple nodes associated with distinct pathways within the same network is more likely to improve response and prevent compensatory signaling across a heterogeneous tumor cell population than targeting several single nodes distributed among different cellular networks (reviewed in^47^). Third, by restricting the extent of inhibition at each single node, we dissipate excess signal flow to avoid activation of compensatory networks that promote resistance and recurrence. The common strategy of maximal inhibition at multiple nodes is more likely to lead to resistance since the surplus may cross the activation threshold of a compensatory driver network. Finally, our goal is to suppress metastasis, the process responsible for cancer lethality, rather than primary tumor growth. By leaving the basic network intact, we are effectively ‘normalizing’ a critical cellular network. The 4D-MAPKi we identified can act as a rheostat, mimicking a metastasis suppressor at low dose but inhibiting growth if higher doses are used. However, by first inhibiting metastasis and associated cellular heterogeneity, we anticipate that even traditional cytotoxic agents will be more effective. As such, this strategy represents a paradigm shift in how we address treatment of metastatic disease in cancer.

## MATERIALS AND METHODS

### Cell lines

MDA-MB-231 (MB231), MDA-MB-436 (MB436), MCF10A, 184A1, and 293T cells were received from American Type Culture Collection (ATCC®). E0771-LMB (LMB) cells were generated by Robin Anderson (Johnstone et al., 2015). M6C cells were generated by Jeffrey Green (Holzer et al., 2003). MDA-MB-231-BM1 (BM1) cells were generated by Massague and colleagues (Kang et al., 2003). SUM159-PT cells were generated by Ethier and colleagues^48^. BM1, MB436, LMB, and M6C cells were cultured in DMEM media with 10% fetal bovine serum (FBS), penicillin (50 U/ml), and streptomycin (50 μg/ml). SUM159-PT cells were cultured in DMEM/F-12 (50/50) with 5% FBS, 5 μg/ml insulin, 1 μg/ml hydrocortisone, and penicillin-streptomycin. MCF10A and 184A1 cells were grown in DMEM/F-12 (50/50) with 10% FBS and penicillin-streptomycin. The cells were used in experiments within 15 passages after their arrival in the laboratory.

### Small molecule inhibitors

For in vitro and in vivo studies, JNK inhibitor SP600125, MEK inhibitor Trametinib (GSK1120212), and MLK inhibitor URMC-099 were purchased from APExBIO (A4604, A3018, B4877, respectively). p38 inhibitor SB203580 was purchased from Selleckchem (Cat No S1076) for the in vitro experiments. For in vivo studies, water soluble SB203580 hydrochloride was purchased from APExBIO (B1285).

### Signaling studies *in vitro*

In order to study the changes in stress kinase signaling upon stress in the presence of RKIP, or small molecule inhibitors of the MAPKs, cells were plated at sub-confluence. Once they reach roughly 70% confluence, they were starved overnight (16-24 hours) in serum-free media, and then induced with anisomycin (Sigma-Aldrich, Cat No A9789) at 25 ng/ml or 100 ng/ml final concentration, 10% serum, or recombinant human epidermal growth factor (EGF) at 10 ng/ml final concentration for 30 minutes to activate MAPK pathways. In studies with small molecule inhibitors of the MAPK pathway, all inhibitors were re-suspended in DMSO and used at indicated concentrations. The cells were pre-treated with the inhibitors in serum-free media for 30 min after overnight serum starvation, immediately before induction with anisomycin, serum, or EGF for 30 minutes. In this case, the inducing agent was directly added to the pre-treatment media that already had the inhibitors, or the pre-treatment mediate was replaced by fresh media containing the inducer and the inhibitors. This is to ensure the inhibitors are present during induction of the MAPK pathways. Upon induction for 30 min, the cells were washed three time with cold PBS and immediately lysed in RIPA buffer for protein collection.

### Protein isolation and western blots

Cultured cells were washed with cold PBS and lysed in RIPA buffer with protease inhibitors (Millipore Sigma, 539134) and phosphatase inhibitors (GoldBio, GB-450). Tumor samples were snap-frozen in liquid nitrogen, pulverized, and lysed in RIPA buffer with protease and phosphatase inhibitors. All samples were sonicated 3 times for 10 seconds at 35% power and centrifuged at max speed for 15 minutes at 4°C. Supernatant was collected and the protein concentration was measured using the Bradford assay. All samples were boiled in 6X Laemmli buffer immediately after protein concentration measurement.

For Western blots, equal amounts of protein, ranging from 10 μg to 50 μg, across all samples were used. Blots were blocked for 1 hour at ambient temperature with either Odyssey® Blocking Buffer (LI-COR Biosciences, 927-40010, diluted 1:1 with PBS) or with 5% FBS in Tris Buffer Saline (TBS) with 0.1% Tween20. Then, blots were incubated with primary antibodies at 4 °C over-night, and with secondary antibodies at ambient temperature for 1 hour. Finally, blots were treated with ECL reagent (Pierce ECL Western Blotting Substrate, Thermo Scientific, 32106) when HRP-conjugated secondary antibodies were used and developed under the Chemiluminescence channel of the LI-COR® Fc Imaging System. Blots with fluorescent secondary antibodies were imaged under 700 nm or 800 nm channels of LI-COR® Fc.

**Primary antibodies used**:

Phospho-RSK2 (Ser227) (Cell Signaling, 3556)
Phospho-ATF2 (Thr71) (Cell Signaling, 9221, no longer available)
Phospho-SEK1/MKK4 (Ser257/Thr261) (Cell Signaling, 9156)
Phospho-TAOK3 (Ser177) + Phospho TAOK2 (Ser181) + Phospho-TAOK1 (Ser181) (Abcam, ab124841)
MLK3 (Cell Signaling, 2817)
Phopsho-p44/42 MAPK (ERK1/2)(Thr202/Tyr204) (Cell Signaling, 9101)
Phospho-MSK1 (Thr581) (Cell Signaling, 9595)
Phospho-SAPK/JNK (Thr183/Tyr185) (Cell Signaling, 9251)
p44/42 MAPK (ERK1/2) (Cell Signaling, 9107)
SAPK/JNK (Cell Signaling, 9252)
Phospho-p38 MAPK (Thr180/Tyr182) (Cell Signaling, 4511)
Phospho-AKT1 (S473) (Cell Signaling, 4060)
Phospho-MKK3 (Ser189)/MKK6 (Ser207) (Cell Signaling, 12280)
Phospho-c-Jun (Ser73) (Cell Signaling, 3270)
Phospho-p70 S6 Kinase (Thr389) (Cell Signaling, 9205)
MLK3 (Abcam, ab51068)
MLK3 (Santa Cruz, sc-166639)
Phopsho-MLK3 (T277/S281) (Abcam, ab191530)
DOCK4 (Santa Cruz, sc-100718)
Casein Kinase 2β (Santa Cruz, sc-12739)
Casein Kinase 2β (Santa Cruz, sc-9030)
TAOK1 (Abcam, ab197891)
TAOK1/PSK2 (Santa Cruz, sc-136094)
TAOK2 (Santa Cruz, sc-47447)
TAOK3 (Abcam, ab150388)
alpha-Tubulin (Santa Cruz, sc-8035)

**Secondary antibodies used**:

Goat anti-Mouse IgG (LI-COR, IRDye® 800CW, 926-32210)
Goat anti-Mouse IgM (LI-COR, IRDye® 800CW, 926-32280)
Goat anti-Rabbit IgG (LI-COR, IRDye® 680RD, 926-68071)
Goat anti-Rabbit IgG, HRP conjugate (EMD Millipore, AP187P)
Goat anti-Mouse IgM, HRP conjugate (Invitrogen, 31440)
Goat anti-Mouse IgG, HRP conjugate (Sigma Aldrich, A4416)

### Transient transfection

Prior to transfection, the cells were plated in 6-well plates and grown to ~70% confluence. siRNA vectors were used at a final concentration of 50nM per well of cells. The vectors were incubated with 10 μl of Lipofectamine 3000 (Invitrogen, L3000-015) in OPTI-MEM media (Gibco, 31985062) for 15-30 minutes. The DNA-lipid complex was then added onto the cells in a drop-wise fashion. Cells were incubated with the siRNAs for at least 24 hours before harvesting for experimental use. All experiments were performed 24-72 hours post-transfection. All siRNA constructs were purchased from Dharmacon:

Individual siGENOME human TAOK1 siRNA, Dharmacon, D-004846-02-0005)
Individual siGENOME human TAOK2 siRNA, Dharmacon, D-004171-13-0005)
Individual siGENOME human TAOK3 siRNA, Dharmacon, D-004844-02-0005)
siGENOME Non-Targeting siRNA Pool #1, Dharmacon, D-001206-13-05)

### Stable Lenti-viral cell line generation

293T cells were plated in T-75 plates and were grown to ~70% confluence prior to transfection. 1 hour prior to transfection, the media was replaced with fresh media. Lentiviral vectors were incubated with 3rd generation viral packaging vectors (pCMV-VSV-G, pMDLg/pRRE, pRSV-Rev) and LT-1 (Mirus, MIR-2305) in OPTI-MEM media for 30 minutes as described by the provider’s instructions. This transfection mix was then added onto the 293T cells in a drop-wise fashion. Virus containing media was collected 24-48 hours after transfection. Cellular content and debris were removed by centrifugation, and the supernatant was filtered through 0.45 μm PES syringe (Millex, SLHP033RS) to remove any remaining cells in the media. Polybrene was added to the media at the final concentration of 8 ng/ml to facilitate viral transduction of the target cell line. The target cell lines were transduced with the virus-containing media for 24-48 hours. At the end of the transduction period, cells were washed, trypsinized, and re-plated for selection. Transduced cells were exposed to high concentration antibiotic selection (3μg/ml puromycin) up to 2 weeks (approximately 3 passages). All lentiviral procedures were carried out following Biosafety Level 3 (BSL3) practices in BSL2 tissue culture hoods according to institutional biosafety rules.

### Boyden Chamber Invasion Assay

Each Boyden chamber membrane (Fisher Scientific, 353097) was coated with a thin layer of BME (200 μl of 0.25 mg/ml stock, or total of 50 μg of BME per membrane) and incubated at 37°C for 1 hour. Cells were trypsinized and centrifuged at 500 x g for 5 minutes followed by two rounds of PBS washes to remove remaining serum-containing media. Then, the cells were resuspended in serum-free media and diluted to the desired concentration for plating onto the Boyden chambers. Each Boyden chamber received 20,000 – 100,000 cells in 300 μl serum-free media, depending on the cell line. 10% serum was used as the chemoattractant for these assays. For the experiments testing the effect of MAPK inhibitors on invasion, the cells were resuspended in drug-containing serum-free media immediately. After 16-24 hours, the membranes were stained with Calcein AM (Fisher Scientific, 354217) for 1 hour at 37°C in the dark to stain for live cells. Cells that are in the top chamber were removed from the membrane with a wet cotton swab. Cells in the bottom chamber were dissociated from the membrane by incubating in cell dissociation buffer (Trevigen, Cultrex® 3455-096-05) in a shaker at 37 °C for 1 hour. Calcein AM signal was measured in Perkin Elmer Victor X3 plate reader as a read-out of invaded cells.

### High-throughput chemotactic invasion assays

For testing anti-invasive drug combinations, IncuCyte® ClearView 96-Well Chemotaxis plates (Essen BioScience) were used. 2,000 cells per well were embedded in 2 mg/ml BME and plated onto the chemotaxis plate following the manufacturer’s instructions. Media containing 2% FBS was used in both top and bottom chambers to maintain cell viability over 72 hours or more. 200 ng/ml human EGF (Bio-Techne, 236-EG-01M) was used as the chemotactic agent in the bottom chamber, and the control wells only had the vehicle for the chemotactic agent.

This assay is more accurate when nuclear-labeled cells are used. Therefore, we generated BM1-mKate2 (nuclear red) cells using IncuCyte® NucLight Red Lentivirus Reagent (Essen BioScience, 4478) following the manufacturer’s instructions. After transduction, cells with the highest nuclear red signal intensity (top 25%) were sorted by FACS.

The chemotaxis module in IncuCyte® can accurately count the number of cells in the top chamber and the bottom chamber of the ClearView plates separately. Invasive capability of the cells in the presence of various small molecule inhibitors was measured as the percentage of cells that moved to the bottom chamber over the period of 72 hours. The formula used for this calculation is (number of cells in the bottom chamber)/(number of cells in the bottom chamber + number of cells in the top chamber) x 100. The total number of cells in the top and bottom chambers is used as a readout of proliferation, which was important for determining drug combinations that blocked invasion without affecting growth properties of the cells.

### Proliferation Assays

For proliferation assays, 1,000 – 20,000 cells (depending on the cell line) were plated in 96-well plates and quantified over 5 days in IncuCyte by measuring confluence in Phase-Contrast images taken every 4 hours. For experiments testing the effect of MAPK inhibitors on proliferation, the cells were plated in 100 μl per well and allowed to adhere overnight. Then, 100 μl growth media containing 2X drug was added directly on top of the initial media.

### 3D Cultures

For 3D proliferation experiments, we used Cultrex® 3D Basement Membrane Matrix, Reduced Growth Factor (Trevigen, 3445-005-01, Lot No 37353J16, Lot concentration: 15.51 mg/ml, referred to as BME). For all experiments, the cells in growth media were mixed with BME at a final concentration of 2 mg/ml. For 3D proliferation assays, 100 μl of the cell/BME mixture was dispensed into each well of a 96-well plate. Upon solidification of BME, 100 μl of growth media was added on top of the solidified gel. For experiments where the cells were treated with inhibitors, the inhibitors were prepared in the growth media at 2X of their desired final concentration and added after the gel is solidified to assure 1X final concentration. The growth of the cells was monitored in IncuCyte® Zoom or S3 models for the indicated duration of time.

### Scratch Wound assays

Migration assays are conducted using IncuCyte “Scratch wound” module. 20,000 – 30,000 cells were plated on IncuCyte® ImageLock Plates (Essen BioScience, 4379). Once the cells reached 100% confluence, the wells were scratched with IncuCyte® WoundMaker (Essen BioScience, 4493) following the provider’s instructions, washed with PBS twice, and supplied with fresh growth media. For experiments testing the effect of MAPK inhibitors on cell migration, fresh growth media containing the inhibitors at the desired final concentration was added onto the cells after the PBS washes. Wound-healing process was monitored over 72 hours in IncuCyte, and wound density was measured over time as a readout of cell migration.

### RNA isolation and qRT-PCR

Cells were washed with cold PBS twice and lysed in TRI Reagent (Zymo Research, R2050-1-200). RNA was isolated using Direct-zol™ RNA MiniPrep (Zymo Research, R2052). 4 μg of total RNA from each sample was converted to cDNA using High Capacity cDNA Reverse Transcription Kit (Applied Biosystems, 4368813). Primer pairs used for this study are listed below.

List of Human primers used in this study:

Hs_PEBP1 Forward: GCTCTACACCTTGGTCCTGACA Reverse: AATCGGAGAGGACTGTGCCACT
Hs_NFATC2 Forward: GATAGTGGGCAACACCAAAGTCC Reverse: TCTCGCCTTTCCGCAGCTCAAT
Hs_ROCK1 Forward: GAAACAGTGTTCCATGCTAGACG Reverse: GCCGCTTATTTGATTCCTGCTCC
Hs_ROCK2 Forward: TGCGGTCACAACTCCAAGCCTT Reverse: CGTACAGGCAATGAAAGCCATCC
Hs_ADAM10 Forward: GAGGAGTGTACGTGTGCCAGTT Reverse: GACCACTGAAGTGCCTACTCCA
Hs_ADAM17 Forward: AACAGCGACTGCACGTTGAAGG Reverse: CTGTGCAGTAGGACACGCCTTT
Hs_EPC1 Forward: CCAGACATGCAGTACCTCTACG Reverse: GCTGTTTCTGCATGAGTGCCAG
Hs_PIKFYVE Forward: CTGAGTGATGCTGTGTGGTCAAC Reverse: CAAGGACTGACACAGGCACTAG
Hs_DOCK4 Forward: GCATGTGGATGATTCCCTGCAG Reverse: GGAGGTGATGTAACACGACAGG
Hs_DOCK5 Forward: GCTTCTGAGCAACATCCTGGAG Reverse: TCCTTCTCAGCAGCCGTTCCAT
Hs_ARL13B Forward: GAACCAGTGGTCTGGCTGAGTT Reverse: GTTTCAGGTGGCAGCCATCACT
Hs_DDR2 Forward: AACGAGAGTGCCACCAATGGCT Reverse: ACTCACTGGCTTCAGAGCGGAA
Hs_ITGA1 Forward: CCGAAGAGGTACTTGTTGCAGC Reverse: GGCTTCCGTGAATGCCTCCTTT
Hs_RAPGEF2 Forward: CTCGGATCAGTATCTTGCCACAG Reverse: AGGTTCCACTGACAGGCAATGC
Hs_RAPGEF6 Forward: AGACAGATGAGGAGAAGTTCCAG Reverse: GACCTCATAGGCACTGGAGACA
Hs_APC Forward: AGGCTGCATGAGAGCACTTGTG Reverse: CACACTTCCAACTTCTCGCAACG

List of Mouse primers used in this study:

Mm_PEBP1 Forward: ACTCTACACCCTGGTCCTCACA Reverse: TGAGAGGACAGTGCCACTGCTA
Mm_NFATC2 Forward: ACTTCACAGCGGAGTCCAAGGT Reverse: GGATGTGCTTGTTCCGATACTCG
Mm_ROCK1 Forward: CACGCCTAACTGACAAGCACCA Reverse: CAGGTCAACATCTAGCATGGAAC
Mm_ROCK2 Forward: GTGACCTCAAACAGTCTCAGCAG Reverse: GACAACGCTTCTGAGTTTCCTGC
Mm_ADAM10 Forward: TGCACCTGTGCCAGCTCTGATG Reverse: GATAGTCCGACCACTGAACTGC
Mm_ADAM17 Forward: TGTGAGCGGTGACCACGAGAAT Reverse: TTCATCCACCCTGGAGTTGCCA
Mm_EPC1 Forward: CTGCCAGGCTTCAGTGCTAAAG Reverse: ACTGACAGCCTGCTTTCCTACG
Mm_PIKFYVE Forward: TCTTCTGCCCAGTCCAGCAATG Reverse: ACAGAACATGCTCGGACACTGG
Mm_DOCK4 Forward: GATAGGAGAGGTGGATGGCAAG Reverse: CGCCTTGAGATGCAGATCGTAG
Mm_DOCK5 Forward: GAGCCGACAGTCTCCTCACATT Reverse: CTGCCTGGTTTTGAAGGTGCTG
Mm_ARL13B Forward: ACCAGTGGTCTGGCTGAGATTG Reverse: CATCACTGTCCTTCTCCACGGT
Mm_DDR2 Forward: TCATCCTGTGGAGGCAGTTCTG Reverse: CTGTTCACTTGGTGATGAGGAGC
Mm_ITGA1 Forward: GGCAGTGGCAAGACCATAAGGA Reverse: CATCTCTCCGTGGATAGACTGG
Mm_RAPGEF2 Forward: GCCGAATGGCATCAGTCAACATG Reverse: CAACATCCAGCACTGTGGCGTT
Mm_RAPGEF6 Forward: ACAGAGTGAGCCAGGTGCTTCA Reverse: CACTCACTTCCTCAGTTGGTCC
Mm_APC Forward: GTGGACTGTGAGATGTATGGGC Reverse: CACAAGTGCTCTCATGCAGCCT

### ChIP assays

BM1 cells were crosslinked with 10% formaldehyde for 10 min and quenched with 0.125 mM glycine for 3 min. Cell were then lysed for sonication at 80% output for 4 times for 10 seconds with a 10 second pause in between each cycle. The lysate was pre-cleared with IgG (Santa Cruz, sc-2028) for 1 hour at 4°C and the supernatant was precipitated with antibodies against BACH1 (AF5776, R&D System), or IgG (normal mouse IgG, Santa Cruz, sc-2025) overnight at 4°C. Primers for ChIP quantitative RT-PCR are listed below.

List of Human primers used for the ChIP assay quantitative RT-PCR

Hs_ROCK1 Forward: CAGCCTCACTCTCCCATTTT Reverse: TCCAGCCTTTCCTCTGCTAA
Hs_PIKFYVE Forward: CTGGACTCCTTCTGCCTGAG Reverse: AAGACTCCGCCCTCTGTTTT
Hs_DOCK4 (upstream) Forward: ATTTGCCTGGAGTGGAAGTG Reverse: CTGTATCCAGGGGGATGATG
Hs_DOCK4 (downstream) Forward: TAAGCCCTAGCTCCTGGACA Reverse: AGGGGTCACAAACACTCCTG
Hs_RAPGEF2 Forward: AAAAATGCCAAGAAGGGGTTA Reverse: CACTCATCTAGACAGACCCCTGA
Hs_RAPGEF6 Forward: CGCCACAGTTCATTCACACT Reverse: GCGAAGGGTTGTTTGCTAGA

### Mouse Studies

Mice were procured and housed by the Animal Resources Center and handled according to the Institutional Animal Care and Use Committee at the University of Chicago. Athymic nude mice were purchased from Harlan Sprague Dawley and C57Bl/6 mice were purchased from the Jackson Laboratories. C3-1-TAg-REAR mice were received from Jeffrey Green and maintained in-house.

For primary tumor growth experiments, 2 × 10^6^ BM1 cells, 5 × 10^5^ LMB cells, or 1 × 10^6^ M6C cells were injected orthotopically near the mammary fat pad of athymic nude, C57Bl/6, or C3-1-TAg-REAR mice, respectively. Tumor growth was monitored over time by caliper measurements of the width and length of tumors. Tumor volumes were calculated with the formula:

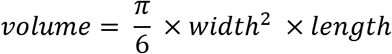

The mice were sacrificed when the tumors reached approximately 1cm^3^.

For metastasis assays, 1×10^5^ luciferase-expressing BM1 cells (BM1-luc) were injected into the left ventricle of the heart to allow for systemic distribution of the bone-tropic tumor cells. 5×10^5^ LMB cells and 1×10^6^ M6C cells were injected into the tail vein. Mice were monitored for 3-6 weeks (depending on the model) for tumor development. At the earliest sign of respiratory problems or paralysis of the limbs, the experiment was ended, and the mice were euthanized. Tumor burden was measured at the end of the study via Xenogen IVIS® 200 Imaging System (PerkinElmer) for BM1-luc tumors. For LMB and M6C tumors, tumor burden was measured by counting overt surface metastases in the lungs after perfusion and formalin fixation, as well as counting tumors in cross-sections of the lungs after H&E staining (described below).

For the *in vivo* studies involving MAPK inhibitors and the 4-drug MAPKi combination treatment, small molecule inhibitors were resuspended under sterile conditions. Since not all of the inhibitors were water-soluble, all inhibitors were initially resuspended in DMSO at the volumes that will result in less than 5% final DMSO concentration. p38 inhibitor SB203580 and MLK inhibitor URMC-099 were further diluted to the desired concentration with 50 %PEG-400 (Sigma, 91893) + 50 %saline. JNK inhibitor SP600125 and MEK inhibitor Trametinib were diluted in corn oil (Sigma, C8267). For the 4-drug combinatorial treatment, all inhibitors were dissolved in their own solvent at 4X higher concentration then the desired final concentration. Then, SB203580 and URMC-099 were mixed at a 1:1 ratio, reducing the concentration for each drug down to 2X. Similarly, SP600125 and Trametinib were mixed at a 1:1 ratio. These dual combination solutions were then filtered through 1.1 μm PES filter syringes to assure sterility. Each mouse received 50μl of each dual combination on the same day, resulting in a total of 100μl of drug mix (2 injections per mouse) with each drug at their desired 1X final concentration. Final concentration for SB203580, URMC-099, SP600125, or Trametinib in the 4-drug MAPKi combination was 10 mg/kg/day, 10 mg/kg/day, 10 mg/kg/day, or 0.5 mg/kg/day, respectively. All injections were intraperitoneal.

For the tumor growth experiments with the MAPK inhibitors, tumors were allowed to reach the size 50-100 mm^3^ size before the MAPKi treatment began. Then, the mice were treated with the respective MAPKi treatment (or the control) for up to 3 weeks. Tumor size was monitored twice a week with a caliper. For the metastasis assays, the tumor cells were treated with the 4-drug MAPKi combination at the in vitro doses for 24 hours prior to injections to allow for anti-metastatic reprogramming of the cells. Homing to metastatic tissues upon intracardiac or tail vein injections can take up to 48 hours. To ensure that the reprogrammed tumor cells do not revert back to their untreated state in the circulation, we pre-treated the mice with the MAPKi combination 2-6 hours before tumor cell inoculation as well. After the inoculation, the mice were treated with the inhibitors daily for up to 3 weeks until the experimental endpoints discussed above were reached.

### Histology

Tumor tissues were fixed in 10% formalin upon dissection for 72 hours and then transferred into 70% Ethanol for long-term storage. Mouse lungs were perfused with PBS before formalin fixation step to allow for tissue expansion and high-quality histological analysis. Fixed tissues were embedded in paraffin and sliced into 5μm sections prior to hematoxylin & eosin (H&E) staining. All tissue processing and staining for this body of work was performed by the University of Chicago Human Tissue Resource Center. For the detection of tissue morphology as well as tumor populations within the lung, lung sections were deparaffinized, immersed in hematoxylin, rinsed in warm distilled water, and treated with eosin. Stained slides were scanned at 10X on a Nikon Eclipse Ti2 Inverted Microscope System.

### MIB-MS analysis

Multiplexed inhibitor beads - mass spectrometry analysis on BM1-VC and BM1-RKIP tumors was conducted as previously described (Duncan et al., 2012). Tumors were grown in athymic nude mice as described previously. Once the tumors reached the size of ~300 mm^3^ they were isolated, flash-frozen in liquid nitrogen, and shipped to the Johnson Laboratories in Chapel Hill. Preparation of the lysate for the MIB-MS analysis, and the mass spectrometry were all performed as described by Duncan et al.

### RNA-sequencing

To compare the transcriptomes of metastatic BM1-VC and non-metastatic BM1-RKIP tumors, 1 × 10^6^ cells were injected orthotopically. When tumors reached approximately 200 mm^3^ size (about 3-weeks post inoculation), we harvested the tumors and flash-froze them in liquid nitrogen. Tumor samples were pulverized immediately, and lysed in TRI Reagent® (Zymo Research, R2050-1-200). RNA was extracted using the Direct-zol™RNA MiniPrep Kit (Zymo Research, R2052) following the manufacturer’s instructions under RNAse-free conditions. In order to prevent contamination of the RNA samples by genomic DNA, the samples were treated with DNAse-I (Zymo Research, E1011-A) for 15 min at ambient temperature on the RNA extraction column. Total RNA was eluted in RNAse/DNAse-free water (Zymo Research, W1001-30) and submitted to the University of Chicago Genomics Facility for further analysis.

RNA quality assessment, library preparation, and sequencing of the tumor RNA samples were all performed by the Genomics Facility staff following the facility’s standardized protocols. Quality of the samples were assessed using a Bioanalyzer, and the samples were determined to be of high quality with an average RNA integrity number (RIN) of 8.6. For the RNA-seq analysis we had 7 control tumors and 5 RKIP-overexpressing tumors, so we chose to generate an individual oligo dT selected, mRNA directional library for each tumor sample without any pooling scheme. All 12 samples were run on the same lane in HiSEQ4000 to generate 50 base-pair long single-end reads.

Bioinformatic analysis of the RNA-seq results were all carried out using the web-based bioinformatics platform Galaxy (usegalaxy.org). Raw “*.fastq” files were uploaded to the Galaxy servers via a file transfer protocol (FTP) software. The reads were analyzed for GC content using “FastQC” and trimmed to remove adaptor sequences using “Trim Galore!”. The reads were mapped to the human genome (hg19) using RNA STAR. In all samples, 70-75% of the reads were uniquely mapped. The resulting “*.bam” files were used to count reads per gene with “featureCounts”. Finally, read counts were normalized and analyzed for differential expression between Control and RKIP-overexpressing samples using “DESeq2”. Principle component analysis on the normalized read counts demonstrated two distinct clusters of samples, separated by the RKIP status.

The raw and processed sequence data are deposited to Gene Expression Omnibus (GEO) with the series accession number GSE128983.

### The Cancer Genome Atlas (TCGA) analysis

For the analysis of patient data, normalized RNA-seq results were accessed through the cBioportal data base (www.cbioportal.org) (Gao et al., 2013). For every TCGA cancer type, the provisional data sets were used for analysis (tagged “TCGA, Provisional” on cBioportal). Lists of genes that correlate with RKIP and BACH1 were also downloaded directly from cBioportal, as the data base already has these correlation matrices generated for each TCGA cancer type. Oncotype and expression heatmap plots were directly generated by cBioportal. Prior to generation of these plots, z-score threshold of 0.5 was arbitrarily chosen to classify patients into high versus low expressors for a particular gene of interest. For example, if a patient’s tumor sample has an RKIP expression level that has a z-score higher than 0.5, then the sample was deemed “RKIP-high”, and if the z-score was below −0.5, the sample was deemed “RKIP-low”. If the z-score falls within −0.5 and 0.5, then the sample was considered as “Intermediate”, or “Other”. Both Pearson and Spearman correlations were used in determining gene-gene correlations and a coefficient cut-off of 0.3 was chosen arbitrarily for both correlation metrics.

The clinical metadata regarding the TCGA breast cancer data set was downloaded from cBioportal. The clinical information on breast cancer patients does not contain TNBC status information. So, we assigned TNBC vs. Non-TNBC status to breast cancer samples by considering immunohistochemistry-based assessment of ER, PR, and HER-2 expression. If the sample was negative for all three of these parameters by IHC, the sample was deemed “TNBC” (n=115). Otherwise, the sample was considered as “Non-TNBC”. For the survival analysis, we used “survival” and “survminer” R packages. TCGA breast cancer patients were separated into two groups: target-gene high expressors (TG-high) and target-gene low expressors (TG-low) based on the expression of 5 motility and adhesion genes (ROCK2, DOCK4, ITGA1, APC, and RAPGEF6). If a patient sample falls above the median for all 5 genes, that patient is deemed TG-high. If a patient sample falls below the median for all 5 genes, that patient is deemed TG-low. Survival probability with respect to time (in months) is plotted in a Kaplan-Meier curve and statistical significance was determined using a logrank test.

### Gene set enrichment analyses

Functional gene set enrichment analysis of the differentially expressed genes in the RNA-seq data as well as the genes that correlate with RKIP and BACH1 was performed using the web-based interface of the Metascape software (metascape.org) (Tripathi et al., 2015). For the identification of pathways and processes enriched in the input gene lists, both “Gene Ontology" (GO) and “Kyoto Encyclopedia of Genes and Genomes" (KEGG) categories were considered. A minimum overlap of 5 genes and an enrichment score of 1.5 were chosen as the enrichment parameters. An adjusted p-value cut-off of 0.05 was chosen as the significance threshold.

## NETWORK MODEL

### Model description

We propose a coarse-grained framework for devising a new cancer treatment strategy based on cellular reprogramming. The networks governing the cell dynamics involve a plethora of components interacting in a non-linear fashion, and its quantitative description would require, in principle, the construction of a large system of coupled differential equations. That approach is unfeasible because of a lack of detailed knowledge of the parameters and chemical reactions that they govern. Hence, an alternative approach was used to describe the signal flow within the network which drives the transcription of BACH1 by MAPKs under stress assuming a steady state regime.

We consider that a cancer cell has a multiplicity of hierarchically structured driver networks responsible for activation of the stress response genes that promote metastasis. The BACH1 driver network is a primary absorber of the stress signal that, when compromised, may generate a surplus of kinase signal(s) to activate secondary compensatory networks, enabling redundancy of stress processing and response. Our data indicate that the BACH1 driver network is fully operational in all cell lines and that the stress signal strength can induce saturation of the activity of all components of the network.

We model the driver network considering the stress signal as a steady state flow through the network pathways and the kinase nodes. The nodes of the network are labeled accordingly with the kinase that we posit they represent. The inflow of a node *i* indicates the arrival of the kinase signal(s) into it. The inflow interacts with the kinase node, and the outflow indicates the product of this interaction. Each node has an activity that regulates the inflow of kinase signal(s), and at each node the outflow of products is equal to the inflow of the signal.

The treatment targeting a specific node of the network will reduce its activity and, hence, its capacity for consuming the products (signal) from its upstream node. The surplus of products from the upstream nodes may be either redirected within the network or, in the case of saturation of the crosstalk links, be redirected to activate a secondary compensatory network. In the case of the surplus being greater than the threshold of activation of the compensatory network, an alternative system (driver network) will be turned on and the stress response will still be functional and capable of promoting metastasis.

### Signal Flow description

Figure 6A shows a phenomenological representation of the BACH1 transcription driver network N1 for the BM1 cell line. The nodes containing a kinase are labeled by an index *i* = 2,…,8 while node 1 indicates the splitting of the stress signal and node 9 denotes the induction of BACH1 transcription as generated by the inflow of upstream kinase signals. The inflow of the *i*-th node is denoted by *ϒ*_i_ while its outflow is *Ω*_i_. Since we are considering a conservative network, we have *Ω*_i_ = *ϒ*_i_, for *i* = 1,…,9. The flow capacity of arrow connecting node *i* to *j* is indicated by *c*_ij_ which is a fraction of *ϒ*_i_ when the node has more than one outgoing pathway. Hence, a coefficient *α*_i_ or *A*_i_ besides a pathway denotes the fraction of the inflow going through it such that *α*_1_ + *α*_2_ + *A*_12_ = 1 and α_k_ + *A*_k_ = 1 where *k* = 3,…,6. The absence of those coefficients indicate that the outflow equals the inflow. The intensity of the stress signal is denoted by Σ and the degree of activation of transcription of BACH1 caused by this signal is indicated by ϕ_BACH1_. For the case of maximal stress signaling, we have ϕ_BACH1_ = Σ. The treatment generates a surplus of products of the functional network denoted by E such that the degree of activation of transcription of BACH1 under treatment is ϕ_BACH1_ = Σ − *E*. The degree of activation of transcription of BACH1 can be rewritten as a fraction of maximal stress, ϕ_BACH1_ 1 − ϵ. For simplicity we only represent treatment targeting MEK and its effect on generating RAF surplus.

Let us write the formulae for the input flows in fractions of the stress signal Σ.

A. We have an input signal being split at node 1 such that Υ_1_ = 1 and, since node 1 has three outgoing pathways, we have Ω_1_ = Υ_1_ = α_1_ + α_2_ + *A*_12_ = 1 which implies on: Υ_2_ = α_1_; Υ_5_ = α_2_; Υ_7_ = *A*_12_.
B. Node 2 has one outgoing pathway and generates an outflow Ω_2_ = Υ_2_ that fully inflows node 3, and hence Υ_3_ = Ω_2_ = Ω_3_. Node 5 has two outgoing pathways, hence Ω_5_ = Υ_5_ = α_3_Υ_5_ + *A*_3_Υ_5_ = α_2_α_3_ + α_2_*A*_3_ and, similarly, node 7 outflow obeys Ω_7_ = Υ_7_ = *A*_12_α_4_ + *A*_12_*A*_4_.
C. The inflow of node 8 is Υ_8_ = A_3_Ω_5_ + *A*_4_Ω_7_ such that Ω_8_ = Υ_8_; the inflow of node 4 becomes Υ_4_ = A_5_Ω_8_ + Ω_3_ such that Ω_4_ = Υ_4_; and the inflow of node 6 is Υ_6_ = A_6_Ω_4_ + α_3_Ω_5_ + α_4_Ω_7_ such that Ω_6_ = Υ_6_; Ω_3_, Ω_5_, and Ω_7_ are defined above.
D. The inflow arriving at node 9 is written as Υ_9_ = α_6_Ω_4_ + α_5_Ω_8_ + Ω_6_ such that in the absence of treatment, ϕ_BACH1_ = Υ_9_ = 1 as it can be verified by direct substitution. Indeed, let us consider Ω_8_ = A_3_Ω_5_ + *A*_4_Ω_7_, Ω_4_ = A_5_Ω_8_ + Ω_3_, and Ω_6_ = A_6_Ω_4_ + α_3_Ω_5_ + α_4_Ω_7_, such that Υ_9_ becomes Υ_9_ = (α_6_ + A_6_) Ω_3_ + ((α_6_A_5_ + A_6_A_5_) A_3_ + α_5_A_3_ + α_3_)Ω_5_ + ((α_6_A_5_ + A_6_A_5_)*A*_4_ + α_5_*A*_4_ + α_4_) Ω_7_. Since α_k_ + *A*_*k*_ = 1 for *k* = 3,…,6, we obtain Υ_9_ = Ω_3_ + Ω_5_ + Ω_7_ = Υ_2_ + Υ_3_ + Υ_7_ = 1.

The parameters governing the flow of information through the network were chosen based upon experimental results, such that the values of the parameters α_*k*_ indicate the strength of absorption of the kinase from the upstream node by the current one. For N1 we have:

|  |  |  |  |  |  |
| --- | --- | --- | --- | --- | --- |
| $\alpha_1 = 0.2$ | $\alpha_2 = 0.3$ | $\alpha_3 = 0.7$ | $\alpha_4 = 0.3$ | $\alpha_5 = 0.5$ | $\alpha_6 = 0.5$ |

and the complementary flow parameters are *A*_12_ = 1 − α_1_ − α_2_ and *A*_k_ 1 − α_k_ for *k* 3,…,6. For N2 we eliminate the crosstalk and the parameters become:

|  |  |  |  |
| --- | --- | --- | --- |
| $\alpha_3 = 1$ | $\alpha_4 = 0$ | $\alpha_5 = 1$ | $\alpha_6 = 1$ |

so that *A*_4_ = 1 and *A*_3_ = *A*_5_ = *A*_6_ = 0.

### BACH1 transcription after treatment

We test our approach by analyzing four treatment scenarios using 1) p38i, 2) MEKi, 3) JNKi, and 4) 4D-MAPKi. The reduction in target kinase activity caused by those drugs will be proportional to a function of the drug activity reduction denoted as ξ_p38_, ξ_MEK_, ξ_JNK_, and ξ_4D-MAPK_ = ξ_p38_ + ξ_MEK_ + ξ_JNK_ + ξ_MLK_. That reduction will cause a surplus of kinase signals from the upstream nodes that is proportional to the reduction in the activity of the target node. Hence, the inflow of the target node will be given by 1 − ξ_p38_Υ_8_, 1 − ξ_MEK_Υ_3_, 1 − ξ_JNK_Υ_6_, and 1 − (ξ_p38_Υ_8_ + ξ_MEK_Υ_3_ + ξ_JNK_Υ_6_ + ξ_MLK_Υ_6_), where Υ_*i*_ is the inflow to the target without treatment. The total surplus generated by each single drug treatment is ξ_p38_Υ_8_, ξ_MEK_Υ_3_, ξ_JNK_Υ_6_, while for the 4D-MAPKi we assume additive effect as a first approximation which results in ξ_p38_Υ_8_ + ξ_MEK_Υ_3_ + ξ_JNK_Υ_6_ + ξ_MLK_Υ_6_.

We will indicate the fraction of inhibition caused by a given treatment dose by *x*, since the drug dosage may vary with its specific function. For the drugs targeting nodes *p38, MEK,* and *MLK*, we can represent the reduction in BACH1 transcription effectively as a function of a dose *x* targeting the *i*-th node:

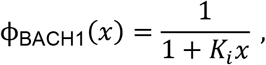

where *K*_i_ is a constant set according to the effect of treatment on reducing BACH1 transcription. The latter is assumed to be equal to the inflow Υ_9_. Then, BACH1 relative transcription after reduction of the activity of node *i* by a fraction ξ_i_Υ_i_ can be written as:

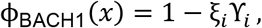

where *i* denotes the node or its corresponding kinase. Hence, we can write the function for the reduction of the node activity as a consequence of drug action as:

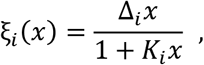

where Δ_i_/*K*_i_/Υ_i_, and Υ_i_ is the inflow at node *i* without treatment.

#### Treatment with p38i

BACH1 transcription after this treatment can be approximated for *K*_1_ = 2, such that:

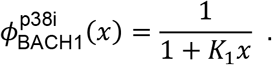

#### Treatment with MEKi

BACH1 transcription after this treatment can be approximated for *K*_2_ = 0.4, such that:

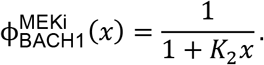

#### Treatment with MLKi

BACH1 transcription after this treatment can be approximated for *K*_4_ = 0.1, such that:

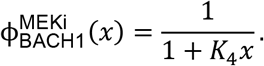

#### Treatment with JNKi

The output of BACH1 after this treatment is obtained using a different model because JNK is repressing nodes 4 and 8. Hence the activity of those nodes will increase because of reduction of activity of node 6. Let us describe the reduction of the activity of node 4 as

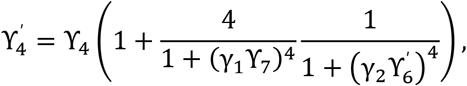

where the primed symbols indicate the inflows under treatment. Since node 4 is also repressed by node 7, and the network is operating under maximal stress, we assume that node 7 represses node 4 maximally such that node 4 activity remains almost constant even under reduction of the activity of node 6. This is based on assuming that the capacity of the pathways connecting nodes 3 and 8 to node 4 will not be affected by a reduction in activity of node 6, and, hence: Υ_4_ = Υ_4_ = *c*_34_ + *c*_84_ = Ω_3_ + *A*_5_Ω_8_.

However, the activity of node 8 will be affected by treatment inactivation of JNK redirecting the flow from nodes 5 and 7. Let us assume that the inflow of node 8 coming from nodes 5 and 7 is, respectively, determined by the activity of node 6 accordingly with

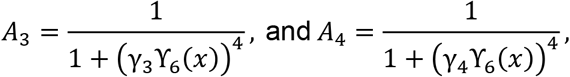

where γ_3_ and γ_4_ are two arbitrary constants and the activity of node 6 can be written as a function of the drug dosage *x* targeting it. The treatment targeting node 6 reduces its activity and induces a compensatory flow to pass through node 8 which is assumed to be capable of absorbing only half of the surplus of kinase signals coming from nodes 5 and 7. Therefore, the other half of surplus will be redirected towards two compensatory networks, each of them receiving the surplus of one of the kinase signals. Then, the surplus is the difference between the absorption by node 8 without and with treatment, and BACH1 transcription becomes

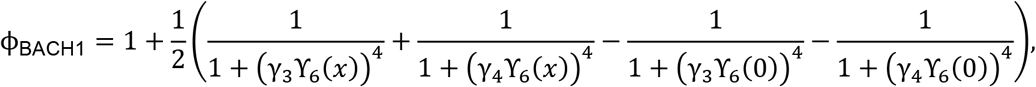

where the factor ½ occurs because we are assuming that node 8 absorbs only half of the surplus generated by treatment. The inactivation of node 6 by treatment can be described by a function

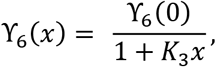

where *K*_3_ = 0.475628 is an arbitrary constant. We also set the values of the constants γ_3_ = 1.520908 and γ_4_ = 1.435780 arbitrarily. Our choice enables us to estimate the node activity under treatment as a fraction of its activity without treatment and to obtain a qualitative description of experimental data based on the reduction of BACH1 transcription following treatment. The analysis of BACH1 transcription after treatment is shown in Figures 6C and 6D for N1 and N2, respectively.

### Surplus analysis

The surplus for each treatment scenario can be evaluated using the output functions after treatment considering that the reduction of activity of a given kinase generates a surplus that comprises the non-absorbed quantities of kinase signal produced at the upstream nodes.

#### Surplus of treatment with MEKi

The total surplus of RAF generated by this treatment can be computed from the transcription of BACH1. The surplus of RAF as function of the treatment is denoted by ϵ_RAF_(*x*), such that ϵ_RAF_(*x*) = ξ_MEK_(*x*)Υ_3_. Since 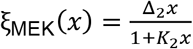, and 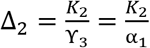, the surplus of RAF is:

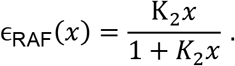

The RAF surplus from MEKi is depicted in Graph 6F, and the function describing it is the same for the 4D-MAPKi treatment. The validity of the formula for both treatment scenarios is because we are assuming that the nodes of the network are operating at their maximal capacity and the surplus is not redirected within the network unless there is a repressive interaction.

#### Surplus of treatment with MLKi

This treatment generates a surplus of stress signal which we assume not being redirected within the driver network. Therefore, this signal will be redirected to a compensatory network and its amount is denoted by ϵ_SIG_(*x*), such that ϵ_SIG_(*x*) = ξ_MLK_(*x*)Υ_5_. Since 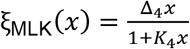, and 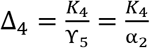, the surplus of stress signal is given by:

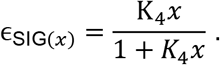

The signal surplus is shown in Graph 6G for the 4D-MAPKi treatment.

#### Surplus of treatment with JNKi and p38i

We compute the surplus generated at nodes 5 and 7 by treatments with JNKi and p38i based on the total surplus that they generate.

#### Total surplus of treatment with JNKi

The surplus generated by this treatment is given by half of the difference between the flow towards node 8 without and with treatment as previously considered when evaluating BACH1 transcription under this treatment. We denote the surplus generated by treatment targeting node 6 as ϵ_6_(*x*) which becomes

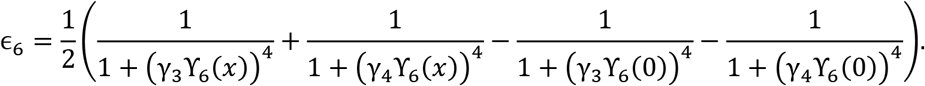

Note, however, that the outflow from node 5 is also affected by treatment by means of its inactivation by drug MLKi. Hence, we have α_2_ → α_2_/(1 + *K*_4_*x*) and the inflow of node 6 becomes

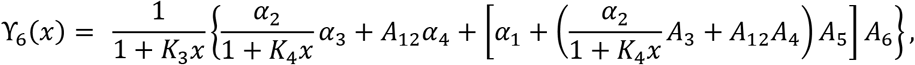

where we are rebalancing the inflow of node 6 without treatment, indicated within curly brackets, by its inactivation because of treatment, which is denoted by the term outside the curly brackets. The term within the curly brackets also has the node 5 outflow reduction because of treatment.

#### Total surplus with treatment with p38i

The surplus generated by this treatment is given by the sum of the inflow towards node 8 without treatment plus the redirected flow from node 6 balanced by the reduction of activity of nodes 5 and 8 because of treatment. We can denote the surplus generated by treatment targeting node 8 by ϵ_8_(*x*) such that

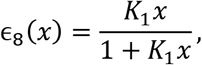

where this surplus is the result of the combination of the surplus signals generated by treatment targeting node 6 and node 8 itself.

The surplus from node 5 can be computed from the sum of the fraction of surplus generated by treatment targeting nodes 6 and 8 where those fractions are proportional to the absorption capacity of each pathway coming from node 5 to nodes 6 or 8. We denote the surplus of node 5 by ϵ_MLK_ such that:

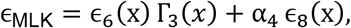

where,

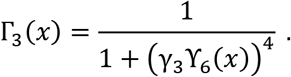

We proceed analogously to compute the surplus of node 7, denoted by ϵ_TAOK_, and obtain:

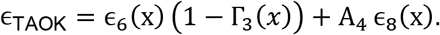

## ACKNOWLEDGMENTS

This study was funded by NIH grants R01 GM121735-01 (to MRR) and CA058223 (to GLJ), the Rustandy fund for Innovative Cancer Research (to MRR), the Women’s Board Scholarship Fund (to AEY) and the Goldblatt Scholarship Fund (to AEY). AFR thanks the Dean of Research of the University of São Paulo for the “Use of Intelligent Systems” call (18.5.245.86.7) and CAPES (88881.062174/2014-01) for financial support. AUS thanks the CAPES foundation for scholarship support. The results shown here are in whole or part based upon data generated by the TCGA Research Network: https://www.cancer.gov/tcga. We thank members of the Rosner Laboratory, Gabor Balazsi, and Robert Rosner for helpful comments.

## AUTHOR CONTRIBUTIONS

Conceptual design of the project; M.R., A.E.Y. Mathematical modeling; A.R., A.S. Mass spectrometry and analysis: T.S., G.L.J.RNA-sequencing data analysis: A.E.Y. Experiments: A.E.Y., D.Y., P.T., J.L., X.X., S.S., C.D., E.S., C.F. Manuscript – writing: M.R., A.E.Y., A.R. Manuscript – review and editing: M.R., A.E.Y., A.R., G.L.J., J.L.

## DECLARATION OF INTEREST

A patent application related to this manuscript, titled “Multidrug targeting of the MAP kinase network to inhibit metastasis”, is pending.

---

**Supplementary Fig. 1:**
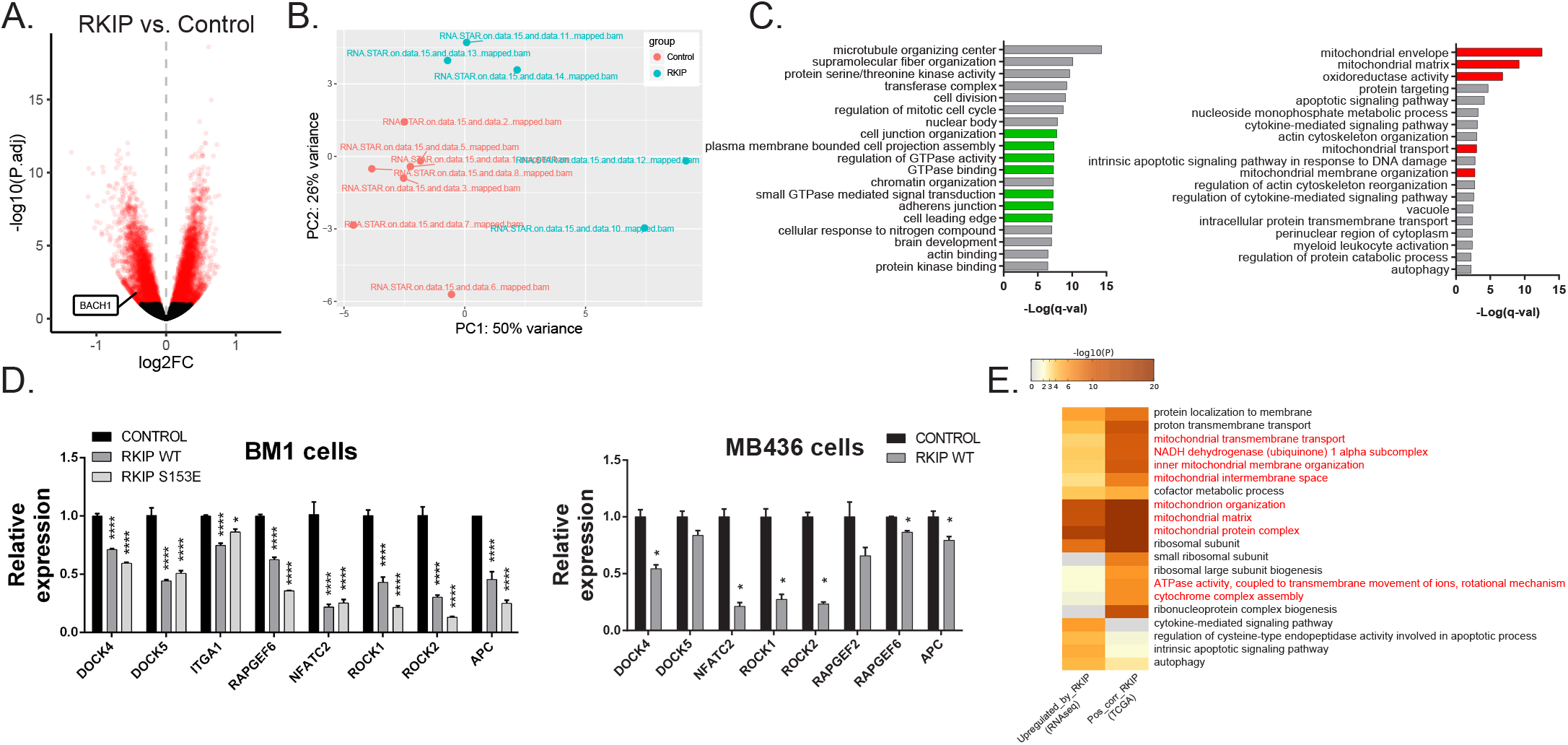
Transcriptional regulation of motility-related genes by RKIP. **A**, Volcano plot highlighting the differentially expressed genes in red, including BACH1, between n=7 control and n=5 RKIP-expressing BM1 tumors. FDR cutoff of 0.1. **B**, Principle component analysis of n=7 BM1-control and n=5 BM1-RKIP tumors used in the RNA-seq analysis. **C**, Gene set enrichment analysis by Metascape of genes downregulated (left) and upregulated (right) by RKIP in the RNA-seq data. FDR corrected p-values are ranked in −log(10) scale. Motility and adhesion related genes are highlighted in green. Mitochondria-related gene sets are highlighted in red. **D**, Regulation of motility gene transcripts by RKIP (wild type or S153E mutant) in BM1 cells (left) and MB436 cells (right) under anisomycin-induced stress conditions in vitro. Data are shown as mean ± s.e.m. of n=3 technical replicates. Statistical significance was determined using one-way ANOVA with multiple testing, with respect to the control sample. **E**, Gene sets commonly enriched in genes positively correlated with RKIP in TCGA BRCA set (provisional set, n=1100), as well as genes upregulated by RKIP in the RNA-seq analysis.

**Supplementary Fig. 2:**
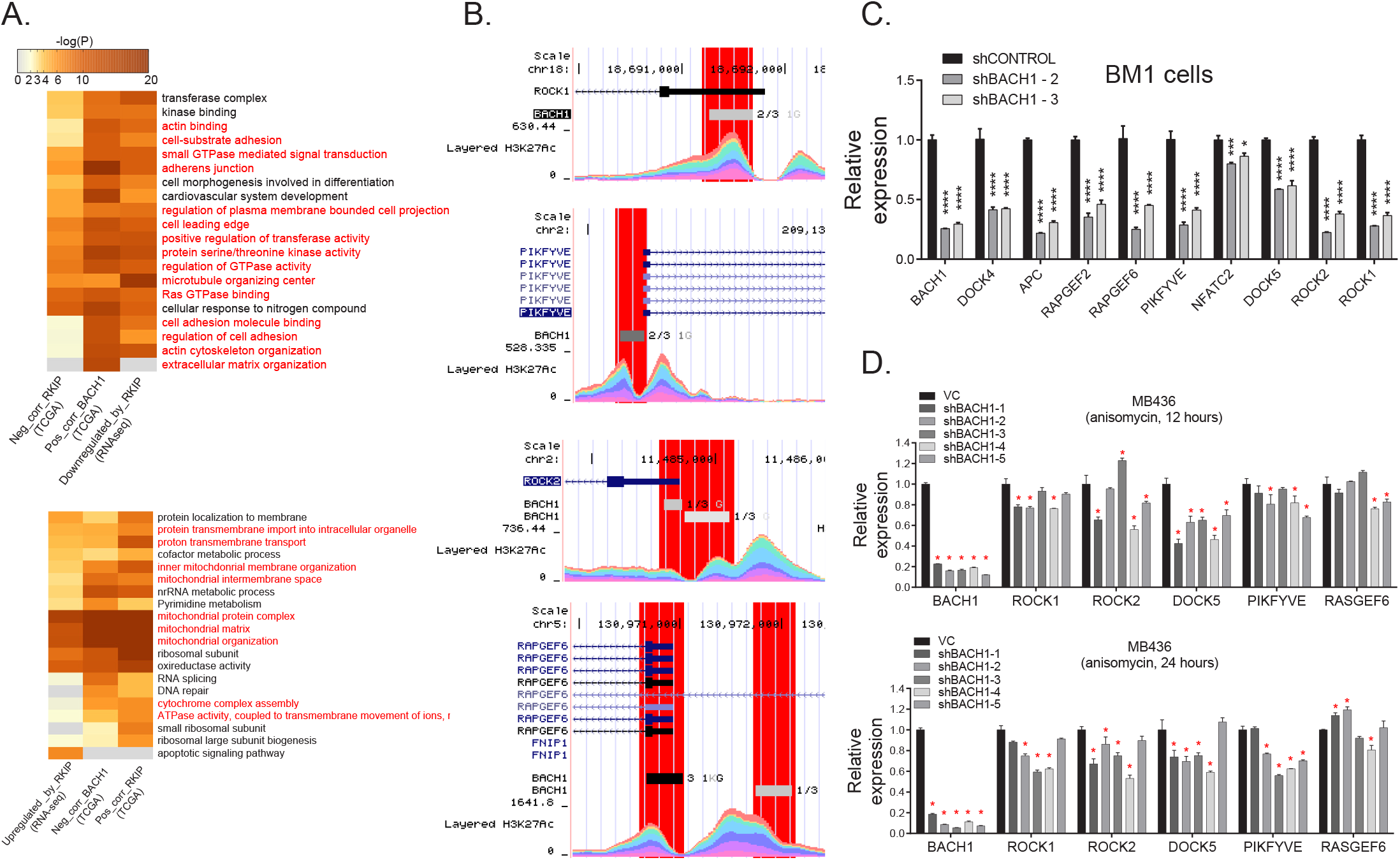
Transcription of metastasis-related RKIP target genes is mediated by BACH1. **A**, Gene sets commonly enriched in genes negatively correlated with RKIP and positively correlated in TCGA BRCA set, as well as genes downregulated by RKIP in the RNA-seq analysis (upper panel). Gene sets commonly enriched in genes positively correlated with RKIP and negetively correlated in TCGA BRCA set, as well as genes upregulated by RKIP in the RNA-seq analysis (lower panel). **B**, BACH1 binding in the promoter region of RKIP target motility genes in ENCODE CHIP-seq data. **C**, Expression of RKIP target motility genes in BACH1-deficient BM1 cells under anisomycin-induced stress conditions. ANOVA with multiple comparisons, *p<0.05, **p<0.01, ***p<0.001, ****p<0.0001 with respect to shCONTROL. **D**, Expression of RKIP target motility genes in BACH1-deficient MB436 cells when induced by anisomycin for 12 hours (upper panel) and 24 hours (lower panel). Student’s t-test, *p<0.05 with respect to the vector control (VC).

**Supplementary Fig. 3:**
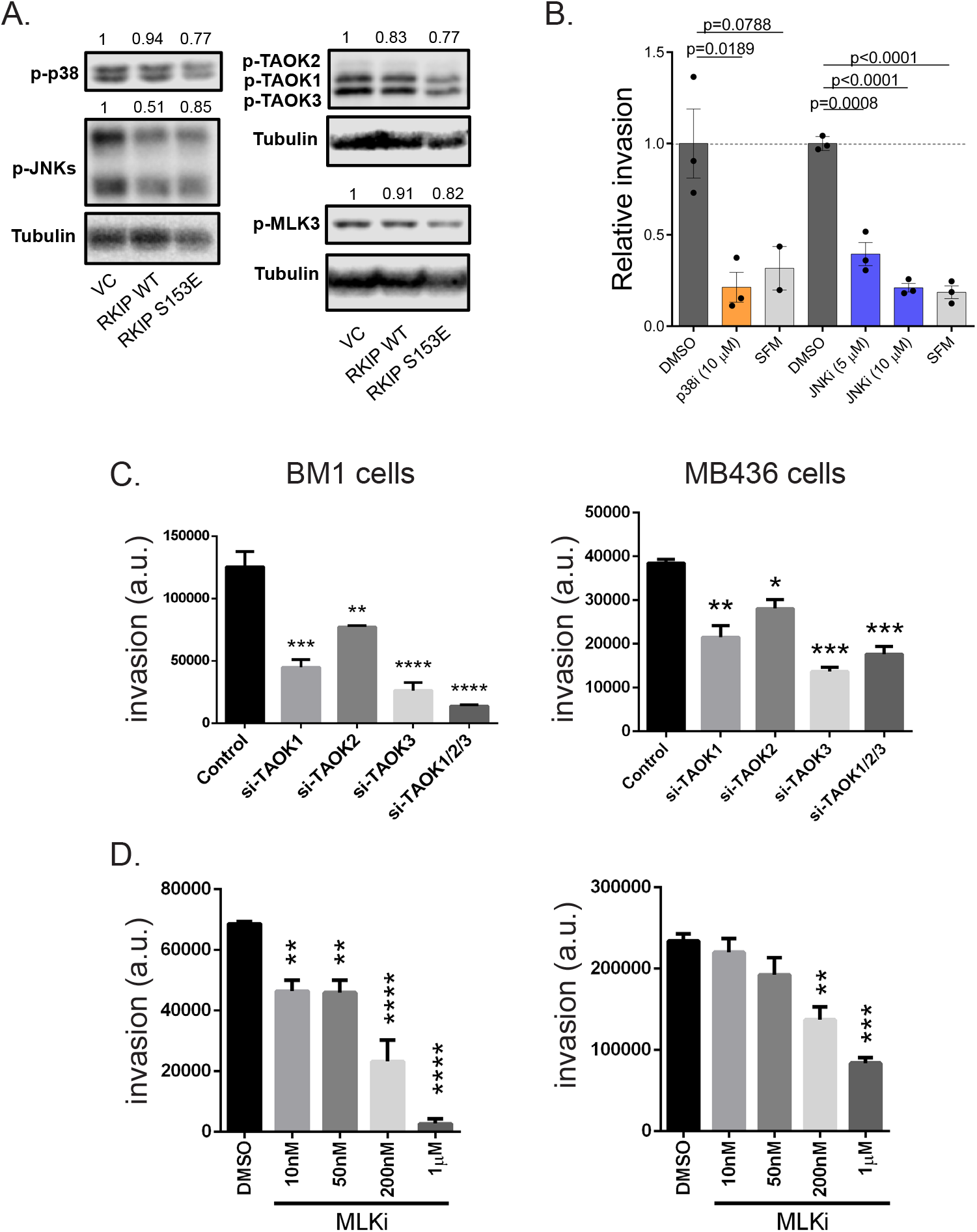
RKIP downregulates the activity of pro-invasive stress MAPKs. **A**, WT or S153E mutant RKIP reduces the activity of stress MAPKs in human TNBC cell line MB436 under anisomy-cin-induced stress conditions in vitro. Quantification for each phospho-kinase band is normalized against the Tubulin signal in the same blot and scaled relative to the empty vector control, which is set to 1. **B**, Invasion of BM1 cells when treated with small molecule inhibitors of p38 and JNK. **C**, Invasion of BM1 cells (left) and MB436 cells (right) when treated with siRNAs targeting TAOKs. **D**, Invasion of BM1 cells (left) and MB436 cells (right) when treated with a small molecule inhibitor of MLK. C and D are representative of two independent experiments. Mean ± s.e.m of technical replicates. Two-tailed t-test with respect to the DMSO-treated or non-targeting siRNA controls. SFM - serum-free media control.

**Supplementary Fig. 4:**
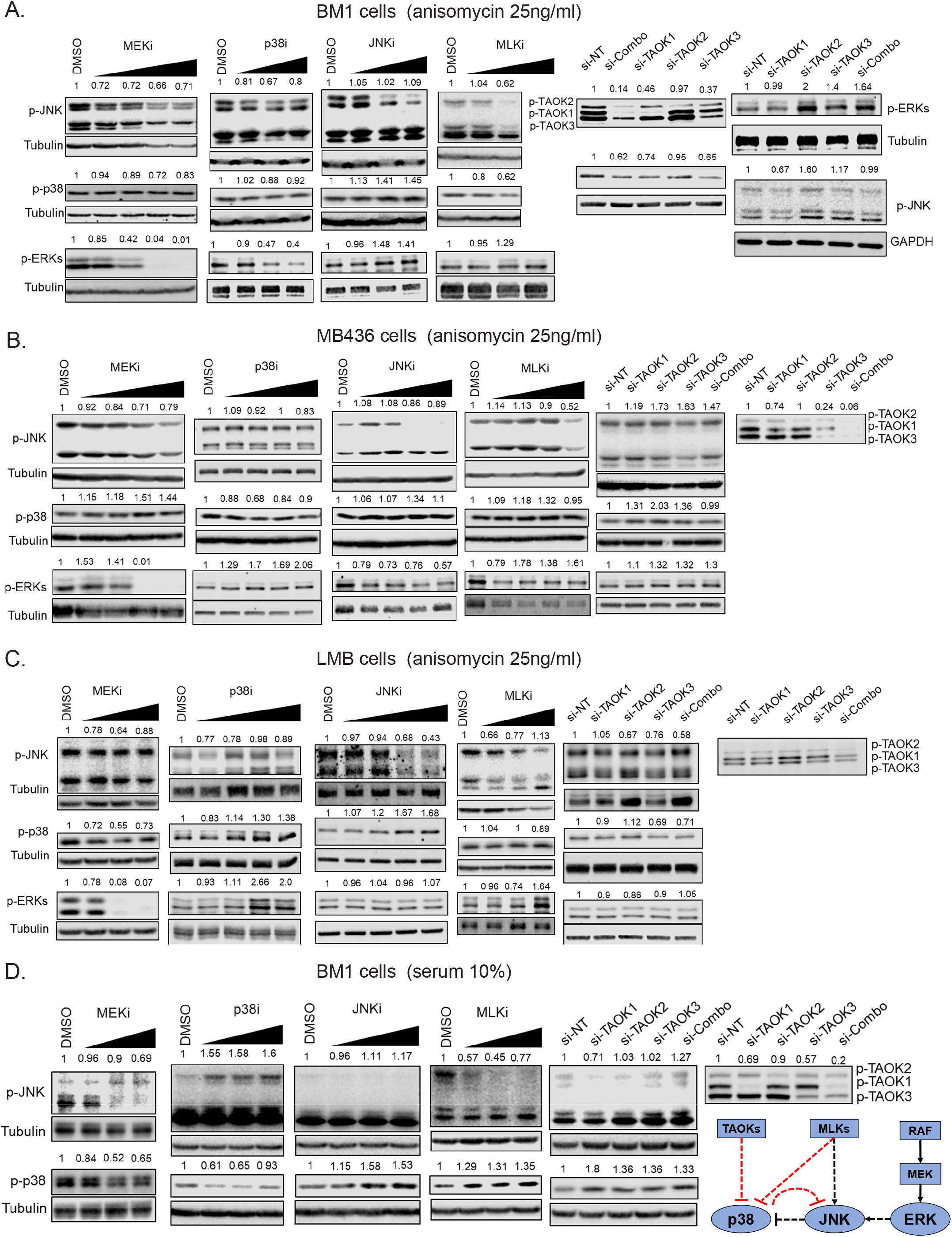
RKIP-regulated MAPK network displays different topologies in a cell type- and inducer-dependent manner. Effect of inhibiting individual MAPKs within the RKIP network on p38, JNK, and ERK activity. BM1 (**A**), MB436 (**B**), and LMB (**C**) cells are induced with anisomycin (25ng/ml) for 30 minutes in the presence of small molecule inhibitors or siRNAs targeting p38, JNK, MEK, MLK, and TAOKs. **D**, p38, JNK, and ERK activity in BM1 cells induced with 10% serum for 30 minutes in the presence of the aforementioned inhibitors. Network diagram highlights the differences in the MAPK network topology when BM1 cells are induced with serum instead of anisomycin (compare to Fig. 2H). All phospho-kinase signal is quantified and normalized to the total tubulin signal in the same sample on the same blot, and scaled to the DMSO-treated control samples.

**Supplementary Fig. 5:**
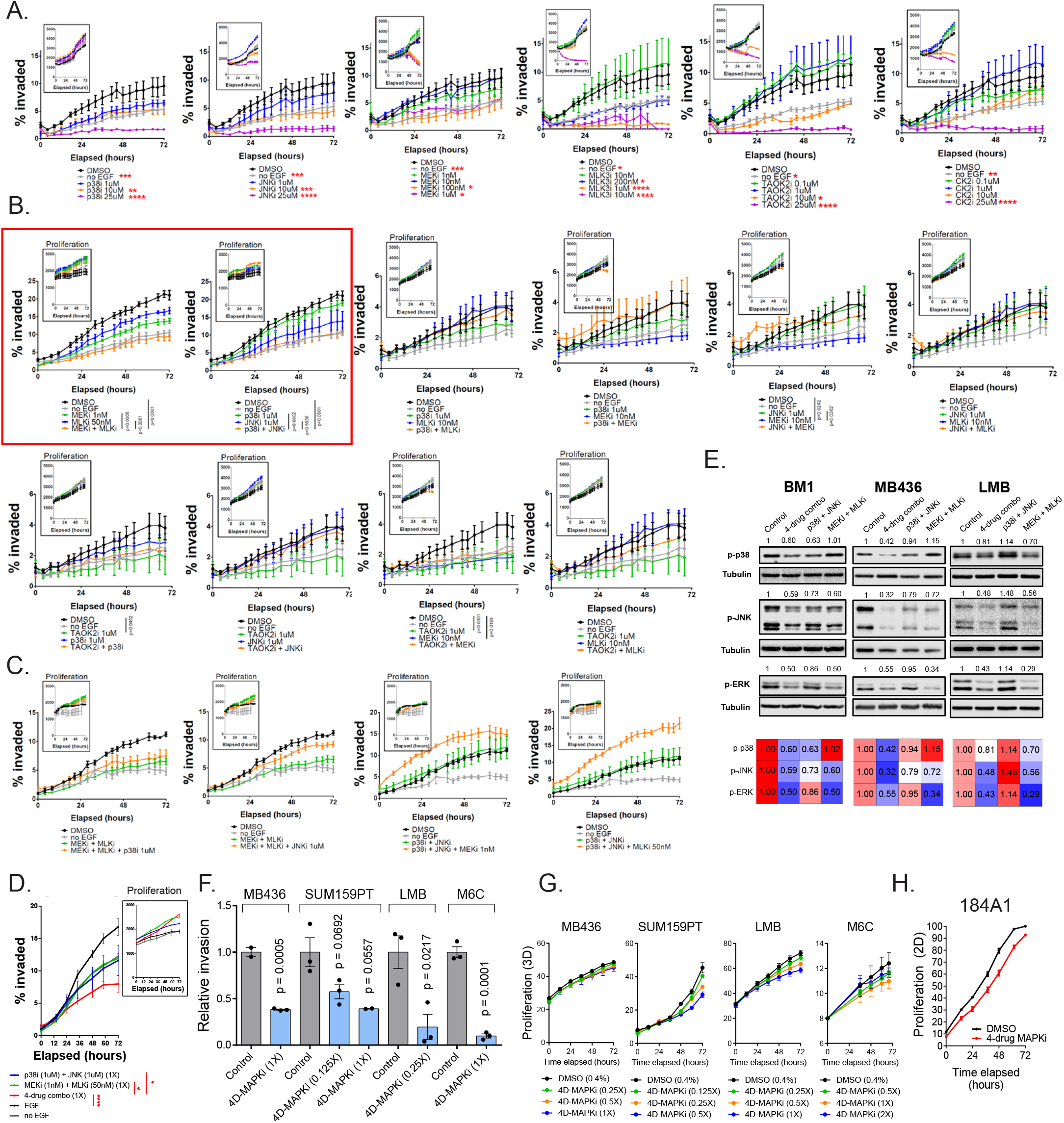
High-throughput invasion assays reveal a low-dose four-drug MAPK inhibitor combination (4D-MAPKi) that phenocopy RKIP in vitro. **A**, Dose escalation experiments with the six inhibitors of the MAPK network demonstrating the range of doses for each inhibitor at which nuclear-labeled BM1 invasion is at least partially reduced, but proliferation is unaffected. Inset shows 3D proliferation of BM1 cells at the same doses over 72 hours. **B**, Dual combinations of MAPK inhibitors tested. The two combinations that showed potential combinatorial effect on BM1 invasion (MEKi+MLKi, p38i+JNKi) are highlighted in a red box. **C**, BM1 invasion and proliferation when a third inhibitor is added to the dual combinations in (B) reveal non-additive nature of MAPK inhibitors used. **D**, Adding MEKi+MLKi and p38i+JNKi into a four-drug mixture has combinatorial effect on invasion. For a-d, mean ± SEM of technical replicates per experimental group. Statistical significance was determined by a two-way ANOVA test with Tukey’s multiple comparison correction. P-values are for the comparisons at 72 hours. **E**, 4D-MAPKi is more effective than the dual combinations in inhibiting all three MAPKs across multiple cell lines under in anisomycin-induced stress conditions. The heatmap shows the densitometry intensity for p-p38, p-JNK, and p-ERKs, normalized the Tubulin signal for each blot and calibrated to the control (DMSO treated) samples. **F and G**, 4D-MAPKi inhibits invasion in a panel of human and mouse TNBC cell lines (F) at non-toxic doses (G). Mean ± s.e.m. of technical replicates per experimental group. Two-tailed students t-test. **H**, 4D-MAPKi is not toxic to normal mammary epithelial 184A1 cells. *p<0.05, **p<0.01, ***p<0.001, ****p<0.0001.

**Supplementary Fig. 6:**
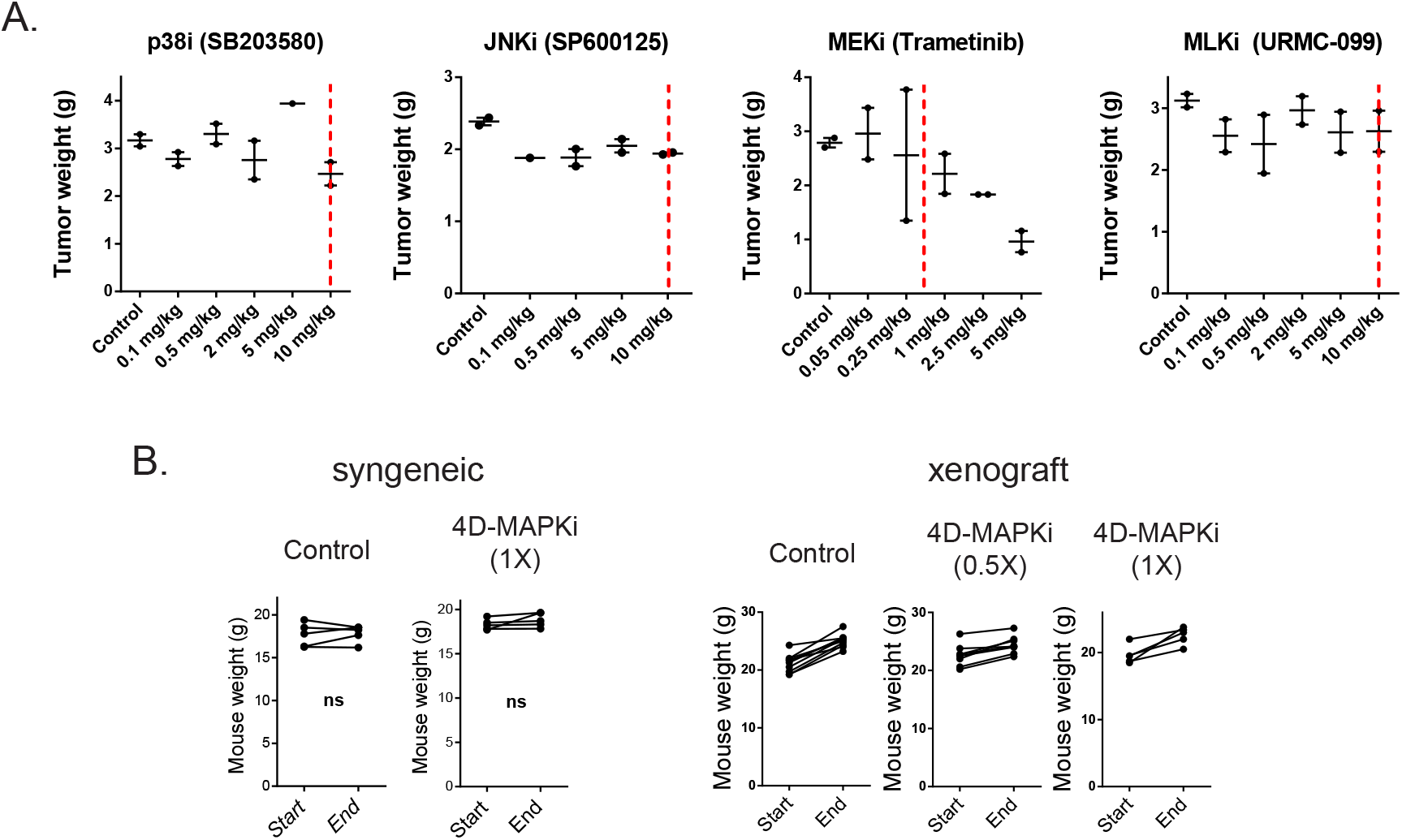
Effect of the individual MAPK inhibitor on tumor growth and their toxicity in combination. **A**, Dose-response experiments reveal that p38i, JNKi, and MLKi do not affect primary LMB tumor growth determined by final tumor weight in a syngeneic model at 10mg/kg/day or lower. MEKi on the other hand, decreases tumor growth in a dose-dependent manner. Red dashed lines indicate the doses chosen to be used in 4-drug combination in vivo. Biological replicates are shown as mean ± s.e.m. **B**, Comparison of mouse weights before and after MAPKi treatment reveal no overall toxicity to the mice due to drug treatment in syngeneic or xenograft models. Paired t-test. ns: not significant.

**Supplementary Fig 7:**
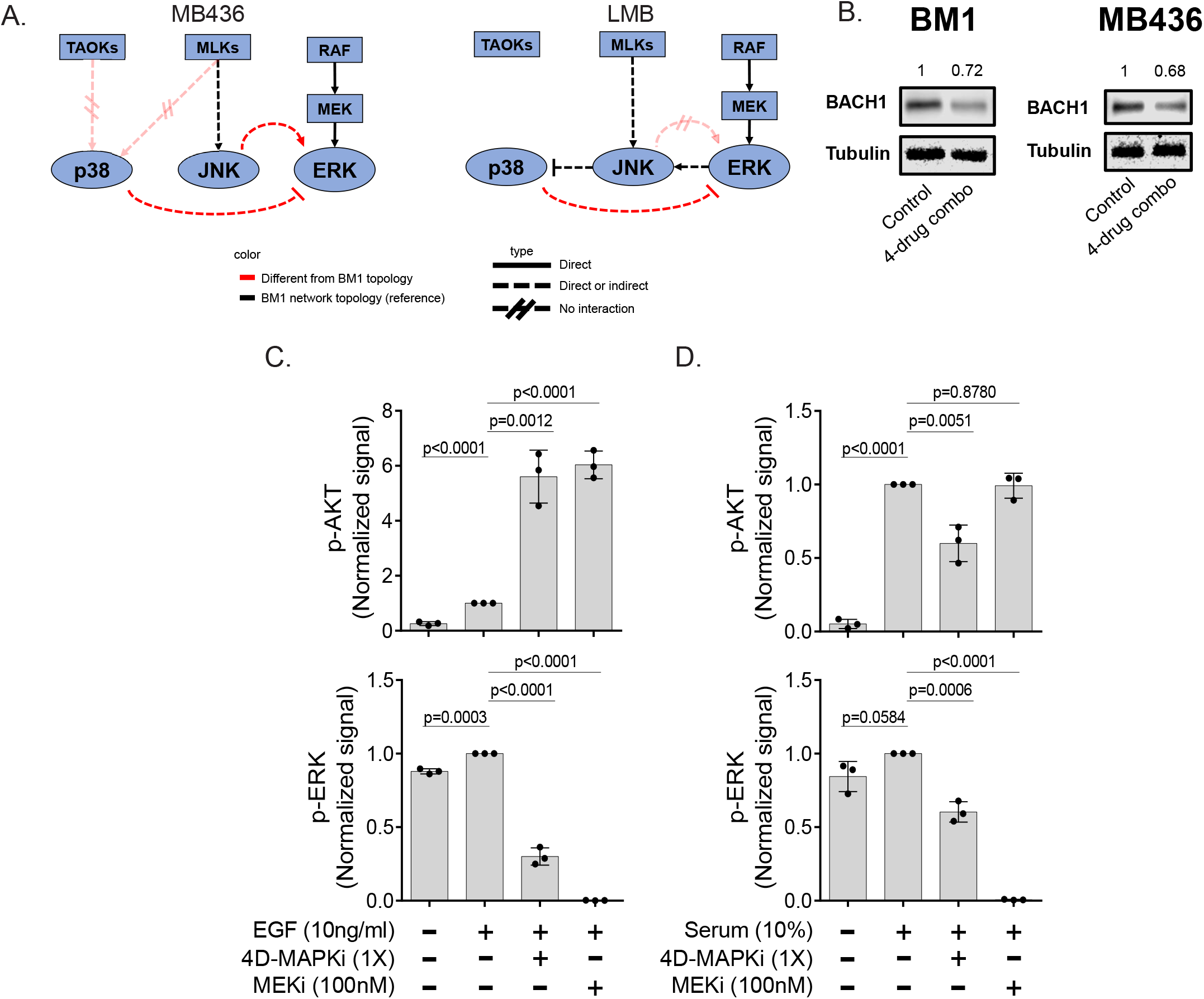
The effect of 4D-MAPKi on the MAPK network output and compensatory network activation. **A**, Topologies of the RKIP-regulated MAPK network in the TNBC cell lines MB436 (left) and LMB (right). Kinase interactions are determined by using small molecule inhibitors or siRNAs against the kinases in the network. Lack of TAOK-p38 interaction was observed in cells treated with a cocktail of siRNAs against all three TAOKs (siCombo). Kinase interactions highlighted in red are the signaling differences between these two cell lines and the reference cell line BM1. **B**, 4D-MAPKi reduces induction of BACH1 protein expression in anisomycin-induced human BM1 and MB436 cells. Western blot images representative of two independent experiments in each cell line. **C and D**, Activation of compensatory AKT signaling in BM1 cells treated with high-dose MEKi or 4D-MAPKi when induced with EGF or serum. Plots demonstrate normalized densitometry analysis of western blots from three independent experiments per induction method.

## Notes

### Competing Interest Statement

The authors have declared no competing interest.

## REFERENCES

1. Weigelt, B., Peterse, J.L. & van’t Veer, L.J. Breast cancer metastasis: markers and models. Nat Rev Cancer 5, 591–602 (2005).

2. Gallaher, J.A., Enriquez-Navas, P.M., Luddy, K.A., Gatenby, R.A. & Anderson, A.R.A. Spatial Heterogeneity and Evolutionary Dynamics Modulate Time to Recurrence in Continuous and Adaptive Cancer Therapies. Cancer Res 78, 2127–2139 (2018).

3. Duncan, J.S. et al. Dynamic reprogramming of the kinome in response to targeted MEK inhibition in triple-negative breast cancer. Cell 149, 307–321 (2012).

4. Wong, C.H., Siah, K.W. & Lo, A.W. Estimation of clinical trial success rates and related parameters. Biostatistics 20, 273–286 (2019).

5. Westin, S.N. et al. Safety lead-in of the MEK inhibitor trametinib in combination with GSK2141795, an AKT inhibitor, in patients with recurrent endometrial cancer: An NRG Oncology/GOG study. Gynecol Oncol 155, 420–428 (2019).

6. Liu, J.F. et al. Results from a single arm, single stage phase II trial of trametinib and GSK2141795 in persistent or recurrent cervical cancer. Gynecol Oncol 154, 95–101 (2019).

7. Robert, C. et al. Five-Year Outcomes with Dabrafenib plus Trametinib in Metastatic Melanoma. N Engl J Med 381, 626–636 (2019).

8. Zhao, M., Li, Z. & Qu, H. An evidence-based knowledgebase of metastasis suppressors to identify key pathways relevant to cancer metastasis. Sci Rep 5, 15478 (2015).

9. Dangi-Garimella, S. et al. Raf kinase inhibitory protein suppresses a metastasis signalling cascade involving LIN28 and let-7. EMBO J 28, 347–358 (2009).

10. Lamiman, K., Keller, J.M., Mizokami, A., Zhang, J. & Keller, E.T. Survey of Raf kinase inhibitor protein (RKIP) in multiple cancer types. Crit Rev Oncog 19, 455–468 (2014).

11. Bonavida, B. et al. Novel therapeutic applications of nitric oxide donors in cancer: roles in chemo- and immunosensitization to apoptosis and inhibition of metastases. Nitric Oxide 19, 152–157 (2008).

12. Chatterjee, D. et al. RKIP sensitizes prostate and breast cancer cells to drug-induced apoptosis. J Biol Chem 279, 17515–17523 (2004).

13. Woods Ignatoski, K.M. et al. Loss of Raf kinase inhibitory protein induces radioresistance in prostate cancer. Int J Radiat Oncol Biol Phys 72, 153–160 (2008).

14. Yun, J. et al. Signalling pathway for RKIP and Let-7 regulates and predicts metastatic breast cancer.EMBO J 30, 4500–4514 (2011).

15. Kang, Y. et al. A multigenic program mediating breast cancer metastasis to bone. Cancer Cell 3, 537–549 (2003).

16. Qin, J.J. et al. NFAT as cancer target: mission possible? Biochim Biophys Acta 1846, 297–311 (2014).

17. Flockhart, R.J., Armstrong, J.L., Reynolds, N.J. & Lovat, P.E. NFAT signalling is a novel target of oncogenic BRAF in metastatic melanoma. Br J Cancer 101, 1448–1455 (2009).

18. Yoeli-Lerner, M. et al. Akt blocks breast cancer cell motility and invasion through the transcription factor NFAT. Mol Cell 20, 539–550 (2005).

19. Amano, M., Nakayama, M. & Kaibuchi, K. Rho-kinase/ROCK: A key regulator of the cytoskeleton and cell polarity. Cytoskeleton (Hoboken) 67, 545–554 (2010).

20. Gadea, G. & Blangy, A. Dock-family exchange factors in cell migration and disease. Eur J Cell Biol 93, 466–477 (2014).

21. Riento, K. & Ridley, A.J. Rocks: multifunctional kinases in cell behaviour. Nat Rev Mol Cell Biol 4, 446–456 (2003).

22. Zhang, Y.L., Wang, R.C., Cheng, K., Ring, B.Z. & Su, L. Roles of Rap1 signaling in tumor cell migration and invasion. Cancer Biol Med 14, 90–99 (2017).

23. Yu, J.R. et al. TGF-beta/Smad signaling through DOCK4 facilitates lung adenocarcinoma metastasis. Genes Dev 29, 250–261 (2015).

24. Kobayashi, M., Harada, K., Negishi, M. & Katoh, H. Dock4 forms a complex with SH3YL1 and regulates cancer cell migration. Cell Signal 26, 1082–1088 (2014).

25. Lee, J. et al. Effective breast cancer combination therapy targeting BACH1 and mitochondrial metabolism. Nature 568, 254–258 (2019).

26. Landt, S.G. et al. ChIP-seq guidelines and practices of the ENCODE and modENCODE consortia. Genome Res 22, 1813–1831 (2012).

27. Yesilkanal, A.E. & Rosner, M.R. Targeting Raf Kinase Inhibitory Protein Regulation and Function. Cancers 10(2018).

28. Yeung, K. et al. Suppression of Raf-1 kinase activity and MAP kinase signalling by RKIP. Nature 401, 173–177 (1999).

29. Tripathi, S. et al. Meta- and Orthogonal Integration of Influenza “OMICs” Data Defines a Role for UBR4 in Virus Budding. Cell Host Microbe 18, 723–735 (2015).

30. Dhillon, A.S., Hagan, S., Rath, O. & Kolch, W. MAP kinase signalling pathways in cancer. Oncogene 26, 3279–3290 (2007).

31. Chen, Z., Hutchison, M. & Cobb, M.H. Isolation of the protein kinase TAO2 and identification of its mitogen-activated protein kinase/extracellular signal-regulated kinase kinase binding domain. J Biol Chem 274, 28803–28807 (1999).

32. Zhou, T. et al. Crystal structure of the TAO2 kinase domain: activation and specificity of a Ste20p MAP3K. Structure 12, 1891–1900 (2004).

33. Cronan, M.R. et al. Defining MAP3 kinases required for MDA-MB-231 cell tumor growth and metastasis. Oncogene 31, 3889–3900 (2012).

34. Isaeva, A.R. & Mitev, V.I. CK2 is acting upstream of MEK3/6 as a part of the signal control of ERK1/2 and p38 MAPK during keratinocytes autocrine differentiation. Z Naturforsch C 66, 83–86 (2011).

35. Sayed, M., Kim, S.O., Salh, B.S., Issinger, O.G. & Pelech, S.L. Stress-induced activation of protein kinase CK2 by direct interaction with p38 mitogen-activated protein kinase. J Biol Chem 275, 16569–16573 (2000).

36. Zhou, B., Ritt, D.A., Morrison, D.K., Der, C.J. & Cox, A.D. Protein Kinase CK2alpha Maintains Extracellular Signal-regulated Kinase (ERK) Activity in a CK2alpha Kinase-independent Manner to Promote Resistance to Inhibitors of RAF and MEK but Not ERK in BRAF Mutant Melanoma. J Biol Chem 291, 17804–17815 (2016).

37. Piala, A.T. et al. Discovery of novel TAOK2 inhibitor scaffolds from high-throughput screening. Bioorganic & medicinal chemistry letters 26, 3923–3927 (2016).

38. Azeloglu, E.U. & Iyengar, R. Signaling networks: information flow, computation, and decision making. Cold Spring Harb Perspect Biol 7, a005934 (2015).

39. Seton-Rogers, S. Signalling: a clearer pathway view. Nat Rev Cancer 14, 156–157 (2014).

40. Mirzoeva, O.K. et al. Basal subtype and MAPK/ERK kinase (MEK)-phosphoinositide 3-kinase feedback signaling determine susceptibility of breast cancer cells to MEK inhibition. Cancer Res 69, 565–572 (2009).

41. Messoussi, A. et al. Recent progress in the design, study, and development of c-Jun N-terminal kinase inhibitors as anticancer agents. Chem Biol 21, 1433–1443 (2014).

42. Okada, M. et al. Repositioning CEP-1347, a chemical agent originally developed for the treatment of Parkinson’s disease, as an anti-cancer stem cell drug. Oncotarget 8, 94872–94882 (2017).

43. Mora Vidal, R. et al. Epidermal Growth Factor Receptor Family Inhibition Identifies P38 Mitogen-activated Protein Kinase as a Potential Therapeutic Target in Bladder Cancer. Urology 112, 225 e221–225 e227 (2018).

44. Cicenas, J. et al. JNK, p38, ERK, and SGK1 Inhibitors in Cancer. Cancers 10(2017).

45. Menzies, A.M. & Long, G.V. Dabrafenib and trametinib, alone and in combination for BRAF-mutant metastatic melanoma. Clin Cancer Res 20, 2035–2043 (2014).

46. Infante, J.R. et al. Safety, pharmacokinetic, pharmacodynamic, and efficacy data for the oral MEK inhibitor trametinib: a phase 1 dose-escalation trial. Lancet Oncol 13, 773–781 (2012).

47. Smith, M.P. & Wellbrock, C. Molecular Pathways: Maintaining MAPK Inhibitor Sensitivity by Targeting Nonmutational Tolerance. Clin Cancer Res 22, 5966–5970 (2016).

48. Forozan, F. et al. Molecular cytogenetic analysis of 11 new breast cancer cell lines. Br J Cancer 81, 1328–1334 (1999).

